# Purification and Characterization of Aspartic Protease Produced from *Aspergillus oryzae* DRDFS13 under Solid-State Fermentation

**DOI:** 10.1101/2020.10.19.346486

**Authors:** Jermen Mamo, Jorge Fernando Suarez Orellana, Vikas Yelemane, Martin Kangwa, Hector Marcelo Fernandez-Lahore, Fassil Assefa

## Abstract

Aspartic proteases (E.C.3.4.23.) are endopeptidases with molecular masses ranging between 30–45 kDa. They depend on aspartic acid residues for their catalytic activity and show maximal activity at low pH. Thus the main objective of the present study was to purify and characterize aspartic protease from locally identified fungi by solid-state fermentation. The aspartic protease in the current study was obtained from *A. oryzae* DRDFS13 under SSF. The crude enzyme extract was purified by size-exclusion (SEC) and ion-exchange (IEC) chromatography. The protein contents of crude enzyme and IEC fractions were determined by BCA methods while the presence of N-glycosylation was checked using Endo-H. Inhibition studies were conducted using protease inhibitors. The milk-clotting activity (MCA), protease activity (PA); molecular weight and enzyme kinetics were determined using standard methods. Optimum temperature and stability, optimum pH and stability, and the effect of cations on MCA were assessed using standard methods. The maximum MCA (477.11 U/mL) was recorded from IEC fraction A_8_. The highest specific activity (183.50 U/mg), purification fold (6.20) and yield (9.2%) were also obtained from the same fraction (IEC A_8_). The molecular weight of 40 kDa was assigned for the purified enzyme (IEC A_8_). However, its molecular weight was decreased to 30 KDa upon deglycosylation assay which infers that the protein is glycosylated. Incubation of the pure enzyme (IEC A_8_) with pepstatin A caused a 94 % inhibition on MCA. The dialyzed enzyme showed a Km and Vmax values of 17.50 mM and 1369 U, respectively. The enzyme showed maximum MCA at 60 °C and pH 5.0 with stability at pH 4.5-6.5 and temperature 35-45 °C. Most cat-ions stimulate the activity of the enzyme; moreover, the highest MCA was detected at 50 mM of MnSO_4_. Furthermore, the results obtained in the present study confirmed that the aspartic protease enzyme produced from *A. oryzae* DRDFS13 and purified in ion-exchange chromatography could be used as a substitute source of rennet enzyme for cheese production.

**Importance:** The production of pure aspartic protease enzyme from local microbes which is useful to substitute shortage of calf rennet enzyme and valuable to diversify cheese production throughout the world.

## 1. Introduction

Aspartic proteases (E.C. 3.4.23.) are endopeptidases having molecular masses between30–45 kDa that depend on aspartic acid residues for their catalytic activity and show maximal activity at low pH. Acid proteases offer a variety of applications in the food and beverage industry and medicine (Vishwanatha, et al., 2009). It plays an important role in the manufacture of cheese in the dairy industry. They can be conveniently produced from different fungal sources using solidstate fermentation and applied in the cheese-making industry owing to their narrow pH and temperature specificities (Rao *et al*., 1998).

*A. oryzae* is one of the best fungi with the highest reported milk-clotting activity (Vishwanatha *et al*., 2009). Optimization of media components such as incubation pH, temperature, and nutrient additions are necessary to fully realize the potential and exploit this species for industrial purposes. These microbial milk-coagulants are also associated with non-specific and heat-stable proteases which lead to the development of bitterness in a cheese after storage and poor yield.

It is, therefore, necessary to purify enzymes with different methods so as to enhance their catalytic activity. The crude enzyme can be easily extracted and purified by a combination of various chromatography and adsorption techniques, that enable adequate separation between the product and other contaminants (Yegin *et al*., 2012).

Glycosylation, a post-translational modification that is found on almost 50% of proteins, in particular, the secreted and transmembrane proteins of eukaryotes, archaea and to a lesser extent in prokaryotes is an important feature of most proteases in higher organisms (Goettig 2016). Glycosylation can potentially affect many proteins’ biochemical properties including, stability, solubility, intracellular trafficking, activity, pharmacokinetics, and antigenicity (Wang et al. 2008). Eukaryotic proteins require glycosylation for proper folding; oligomerization and solubility, as it significantly prolongs the stability and half-life time in many cases by protection against proteolysis (Goettig 2016). Glycosylation has also been known to increase the thermal stability of proteins (Wang *et al*., 2008), as well as against molecular damage by free radicals. Although N-glycosylation is more frequent, O-glycosylation can similarly protect against general and specific proteolysis (Goettig 2016).

Therefore the main objective of the present study was to produce, purify and characterize fungal aspartic protease from the locally identified fungi by solid-state fermentation

## 2. Material and Methods

### 2.1. Medium and cultural conditions for solid-state fermentation for fungi (M2)

The solid-state substrate was prepared in 250 mL Erlenmeyer flask containing 10 g of Durum wheat bran (WB) moistened by adding 12 mL of (0.2M) HCl by mixing thoroughly and autoclaved at 121 °C for 30 min. Then 0.5 mL of 0.5*10^6^ spore suspensions were inoculated into SSF media and incubated at 30 °C for 5 days according to the method by Fernandez-Lahore *et al*., (1998) (Table 1).

**Table 1:**
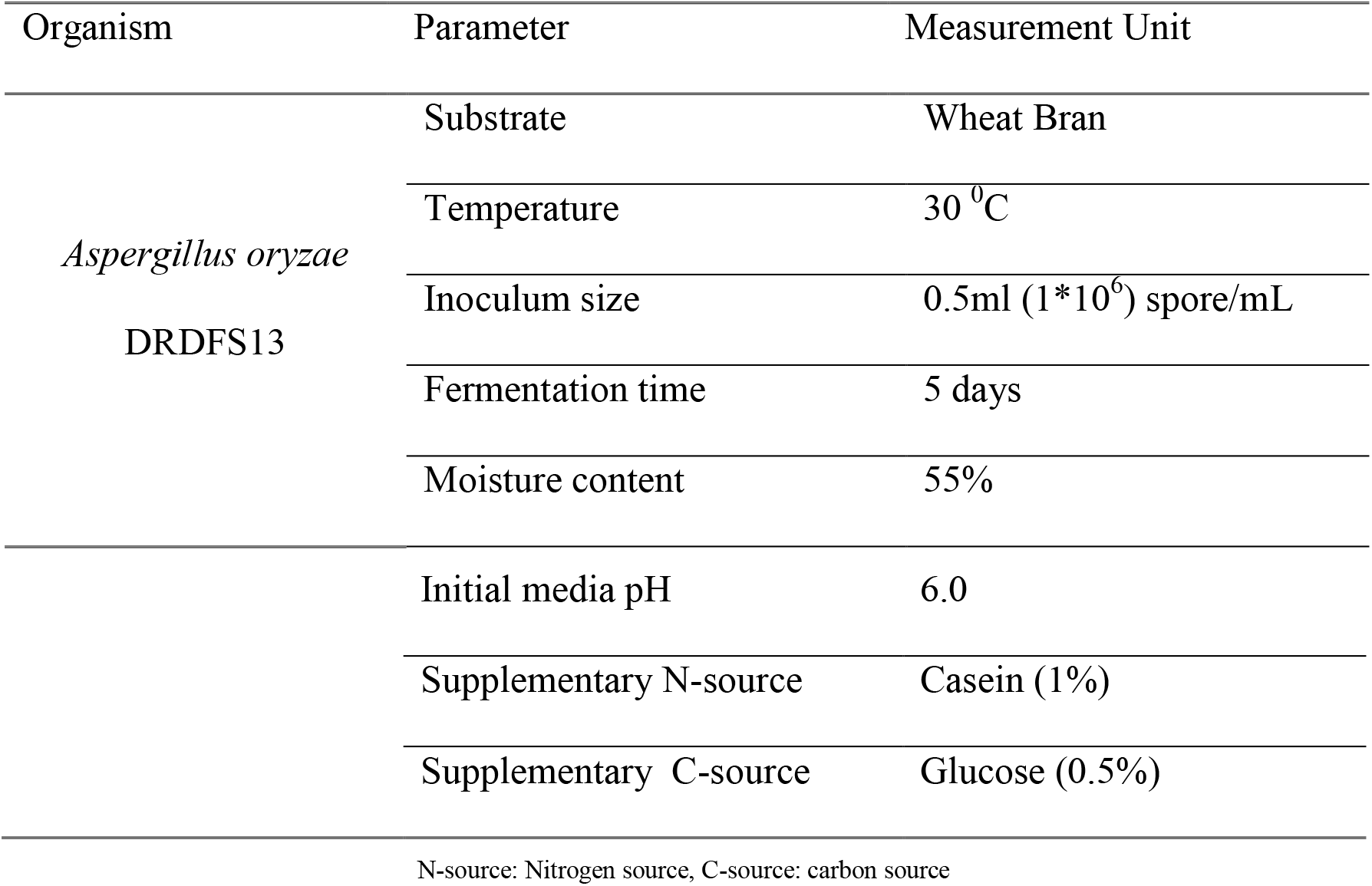
Optimized parameter used for cultivation of *Aspergillus oryzae* DRDFS132.

| Organism | Parameter | Measurement Unit |
| --- | --- | --- |
| *Aspergillus oryzae*<br>DRDFS13 | Substrate | Wheat Bran |
|  | Temperature | 30 °C |
| | Inoculum size | 0.5ml ($1 \times 10^6$) spore/mL |
|  | Fermentation time | 5 days |
|  | Moisture content | 55% |

|  |  |
| --- | --- |
| Initial media pH | 6.0 |
| Supplementary N-source | Casein (1%) |
| Supplementary C-source | Glucose (0.5%) |

N-source: Nitrogen source, C-source: carbon source

### 2.2. Enzyme extraction in solid-state fermentation

The fungal culture was dispensed in 100 mL distilled water (1:10 ratio) and shaken on a rotary shaker (MaxQ 2000 Open-Air Platform Shaker, Thermo Fisher Scientific, USA) at 240 rpm at room temperature. The mixture was centrifuged (Heraeus Pico17/21 37 centrifuge, Thermo Electron Led, Germany) at 10000 rpm at 4 °C for 10 min to recover the supernatant as a crude enzyme.

### 2.3. Assay for milk-clotting activity

The MCA of the enzyme was undertaken according to Arima *et al*., (1970). Accordingly, 0.1 mL of the crude enzyme was added to 1 mL of reconstituted skim-milk (Nestle TM) in a 10 mL test tube pre-incubated at 35 °C for 10 min. Reconstituted skim-milk (NestleTM) solution consisted of 10 g dry skim-milk/100 mL, 0.01 M CaCl2 (AppliChemTM). The appearance of the first clotting flakes was visually evaluated and quantified in terms of Soxhlet units (SU). The endpoint was recorded when discrete particles were visible. The clotting time T (s), the period of time starting from the addition of crude enzyme to the appearance of the first clots and the clotting activity was calculated using the following formula:

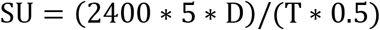

Where T = clotting time (s) D= dilution of crude enzyme

One SU is expressed as the quantity of enzyme required to clot 1 ml of a solution comprising 0.1 g skim milk powder and 0.01M calcium chlorides at 35 °C within 40 min.

### 2.4. Assay of protease activity

The proteolytic activity was assayed according to Arima *et al*., (1970). Accordingly, 0.5 ml of the enzyme extract was added to 2.5 mL of 1% (w/v) soluble casein in 20 mM potassium phosphate buffer at pH 6.5, and the mixture was incubated in a water bath at 35 °C for 10 min. After having added 2.5 mL of 0.44 M trichloroacetic acid to terminate the reaction, the mixture was filtered through Whatman No 1 (90 mm) filter paper. The filtrate was then mixed with 1 mL volume of three times diluted 2N Folin/phenol reagent and 2.5 mL of 0.55 M sodium carbonate solutions and incubated at 35 °C for 20 min to detect color development and measure optical density (OD) on a spectrophotometer (UV-vis, Liantrinsat, and Model-CF728YW-UK) at 660 nm. One unit (1 U) of enzyme activity was defined as the amount of enzyme that liberated 1μg of tyrosine per 1 mL in 1 min.

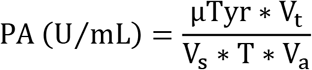

Whereas PA: Protease activity, μTry: μg of tyrosine equivalent released, Vt: Total volume of assay in mL (5 ml of substrate plus 1 ml of Enzyme plus 5 ml of (TCA), Vs: Sample volume (ie. The volume of protease used for assay in mL), T: reaction time (i.e Time of incubation in minutes, 10 min), Va: Volume of assayed (i.e the final volume of the product used in calorimetric determination)

### 2.5. Protein determination

The total protein of the crude extract was determined using Bicinchoninic acid (BCA) methods (Walker, 1996). The working reagent was prepared by mixing 24.5 mL of BCA (Thermo Scientific™) reagent A with 0.5 mL BCA™ Reagent B into 50 mL beaker. The protein content was colorimetrically determined (Eppendorf Biophotometer, Model # 61318, Marshal Scientific, Germany) at 562 nm against a standard prepared from known concentrations of bovine serum albumin (BSA). The protein concentration of the samples (μg/mL) per incubation time was determined using standard.

### 2.6. Crude Enzyme Preparation (Dialysis)

The clarified culture supernatant was subjected to dialysis against 20 mM phosphate buffer (pH 6.0) utilizing a 10 kDa cut-off membrane. After dialysis, a crude enzyme preparation was obtained by concentration by contacting with carboxymethyl cellulose overnight (4 °C). The crude enzyme was resuspended in sodium phosphate buffer (20 mM, pH 6.0) (Ramachandran and Arutselvi, 2013).

### 2.7. Enzyme purification and characterization

#### 2.7.1. Enzyme purification by ion-exchange chromatography

The crude enzyme extract was purified using ion-exchange chromatography (IEC) on a DEAE Sepharose Fast Flow column (5 X 0.5 cm) with an ÄKTA purifier system (GE Healthcare, Munich, Germany). One hundred fifty mL of sample was applied to the column using Buffer A (40 mM citrate-phosphate buffer, pH 6.0) as the equilibration and loading solution, while Buffer B (Buffer A containing 1M sodium chloride) was used as the elution buffer. The purification was run at a flow rate of 2.0 ml per minute and a linear gradient was applied for a period of 5 column volumes. The chromatography fractions (1 mL) were analyzed for enzyme activity and protein content. Fractions showing enzyme activity were pooled and concentrated (Yegin *et al*., 2012).

#### 2.7.2. Size exclusion chromatography

Size-exclusion chromatography was performed with an ÄKTA purifier system (GE Healthcare, Munich, Germany). Hence the crude enzyme extract was 1^st^ centrifuged for 20 min at 16,000 rpm and concentrated using vacuum concentrator. The SEC performed using 2 prepacked columns and the columns were equilibrated using 25 mL a 0.1 M solution of sodium phosphate buffer at pH 6.4 and 2.5 mL of the sample was injected into the column and eluted with 3.5 mL 0.1 M sodium phosphate buffer pH 6.4 (Yegin *et al*., 2012).

#### 2.7.3. Molecular weight determination

The molecular weight of the enzyme was estimated using 12% SDS–PAGE (sodium dodecyl sulfate-polyacrylamide gel electrophoresis) (Yegin *et al*., 2012). The purified enzyme was loaded on gel and electrophoresis was run at 300 V, 35 mA for 55 min. The protein bands were visualized using Coomassie Brilliant Blue solution (AppliChem^™^) staining overnight and destained overnight with destaining solution. The molecular weight of the protein was determined using the prestained protein marker as a standard.

#### 2.7.4. Deglycosylation studies

For deglycosylation assay, 1 mL of the IEC fraction was concentrated for 4 h in vacuum to which 20 μL of the dried sample and 2 μL of 10xGlyco Buffer (New England Biolabs^™^) were added into 2 mL Eppendorf tube and incubated in the thermomixer at 99 °C for 10 min. The content was mixed with 2 μL of denaturing buffer (New England Biolabs^™^) and 2 μL of endo-β-N-acetylglucosaminidase-H (Endo-H, EC 3.2.1.96) (New England Biolabs™) and incubated in the thermomixer at 37 °C for one h. Finally, SDS-PAGE analysis was conducted by loading the mixture of 20 μL the sample (pure sample (non-deglycosylated), deglycosylated sample and pure Endo H) with 20 μL of protein loading dye containing ß-mercaptoethanol. Protein bands were visualized from the gel by staining with Coomassie blue (Appl Chem TM) and destained with the destaining solution (Sigma Aldrich^™^) for one day. The molecular weight was determined from a linear semi-logarithmic plot of relative molecular weight versus the migration distance using standard molecular weight marker protein. The relative mobility (Rf) was also determined as follows.

Rf = migration distance of the protein/migration distance of the dye front

#### 2.7.5. Inhibition study

The IEC fraction A_8_ was subjected to inhibition studies using four protease inhibitors: iodoacetamide, cysteine protease inhibitors (1mM, 10mM); ethylenediaminetetraacetic acid (EDTA) metalloprotease inhibitors (5 mM, 10 mM); phenylmethyl sulphonyl fluoride (PMSF), serine protease inhibitors (1mM,10mM); and pepstatin A, aspartic protease inhibitors (0.02mM, 0.04mM, 0.06mM and 0.08mM) using the method of Yegin *et al*., (2012). The mixtures were incubated at 35 °C for 30 min to determine the residual milk-clotting activity. Residual MCA was defined as the percentage of the activity determined in the absence of inhibitors. The milk-clotting activity and protease activity of the enzyme was also determined according to Arima *et al*., (1970).

#### 2.7.6. Determination of optimum pH and pH stability

The effect of pH on the milk-clotting activity of the enzyme was determined according to Yegin *et al*., (2012). Consequently, 0.01 M CaCl_2_ was prepared in 20 mM citrate-buffer having the pH values (4.5-6.0) and 10 mM phosphate buffer having pH values (6.5-8.0). Ten grams of skim-milk powder dissolved in 100 mL of 0.01 M CaCl_2_ solution that was prepared in 20 mM citrate buffer and 10 mM phosphate buffer and used for the milk-clotting assay. The pH stability was determined by incubating the enzyme sample diluted 10 times in 20 mM citrate buffer having different pH (4.5, 5.0, 5.5 and 6.0) and 10 mM potassium phosphate having pH (6.5, 7.0, 7.5 and 8.0) at 25 °C for 1 h. After incubation, the samples were examined for residual and/or relative milk-clotting activity using the standard procedures (Ageitos *et al*., 2007; Yegin *et al*., 2012).

#### 2.7.7. Determination of optimum temperature and temperature stability

For the determination of the optimum temperature of the enzyme, the milk-clotting activity of the reaction mixture (10 % skim-milk (w/v) in 0.01 M CaCl_2_) was assayed at different temperature ranging from 25-70 °C with 5 °C interval. Likewise, the thermal stability was determined by incubating the enzyme at a temperature ranging from 35-60 °C with with 5 °C interval for 15 and 30 min. After incubation, the enzyme was cooled in an ice bath and residual milk-clotting activity was determined as described before (Yegin *et al*., 2012).

#### 2.7.8. Effect of substrate concentration on the milk-clotting activity

To determine the effect of substrate concentrations on the milk-clotting activity of the enzyme, 25, 50, 100, 150 and 200 g/L of the skim-milk were taken and tested under the same reaction mixture and conditions as before (El-tanboly *et al*., 2013).

#### 2.7.9. Effect of metal ions on the milk-clotting activity

The effect of varying monovalent and divalent metal ions on the milk-clotting activity in the forms of chloride and sulfate salts (Na, K, Ca^2+^ Mg^2+^ Mn^2+^, Fe^2+^, Zn^2+^, Ni^2+^, Cu^2+^ and Co^2+)^ were added to the substrate at a final concentration of 10 mM and the milk-clotting assay was conducted after 30 min (Table 2). For comparison, the control (without the addition of CaCl_2_ or any metal ion) was taken as 100% and the effect of various metal ions was expressed as residual or relative milk-clotting activity (Kumari *et al*., 2016).

**Table 2:** Metals ions used for milk-clotting assay

| S.No | Type of Metal Ions | Concentrations |
| --- | --- | --- |
| 1. | Control (without adding any metal ion) | - |
| 2. | $\text{NaCl}$ | 10 mM |
| 3. | $\text{KCl}$ | 10 mM |
| 4. | $\text{CaCl}_2$ | 10 mM |
| 5. | $\text{MgCl}_2$ | 10 mM |
| 6. | $\text{FeCl}_2$ | 10 mM |
| 7. | $\text{MnCl}_2$ | 10 mM |
| 8. | $\text{NiCl}_2$ | 10 mM |
| 9. | $\text{MgSO}_4 \cdot 7\text{H}_2\text{O}$ | 10 mM |
| 10. | $\text{MnSO}_4 \cdot 7\text{H}_2\text{O}$ | 10 mM |
| 11. | $\text{ZnSO}_4 \cdot 7\text{H}_2\text{O}$ | 10 mM |
| 12. | $\text{CuSO}_4 \cdot 7\text{H}_2\text{O}$ | 10 mM |
| 13. | FeSO<sub>4</sub>.7H<sub>2</sub>O | 10 mM |
| 14. | CoSO<sub>4</sub> | 10 mM |

#### 2.7.10. Effect of MnSO_4_ and CaCl_2_ concentration on the milk-clotting activity

To study the effect of MnSO_4_ and CaCl_2_ on the clotting efficiency of the dialyzed enzyme, various concentrations of MnSO_4_ and CaCl_2_ (0.00 M, 0.005 M, 0.01M, 0.05M, 0.1M, and 0.2M) were incorporated in the reaction mixture (Table 3). Time taken for the appearance of the first clot was recorded and compared with the control sample (El-Tanboly *et al*., 2013).

**Table 3.** Concentration of MnSO_4_ and CaCl_2_ used for milk-clotting assay

| S.No | Conc. Of MnSO<sub>4</sub> in mM | Conc. of CaCl<sub>2</sub> in mM |
| --- | --- | --- |
| 1 | 0.00 | 0.00 |
| 1. | 5.00 | 5.00 |
| 2. | 10.00 | 10.00 |
| 3. | 50.00 | 50.00 |
| 4. | 100.00 | 100.00 |
| 5. | 200.00 | 200.00 |

#### 2.7.11. Michaelis-Menten constant

The proteolytic activity was assayed on casein solutions at a concentration in the range of 0 - 2% in 20 mM potassium phosphate buffer (pH 6.5) (Otani *et al*., 1991). The kinetic parameter (Vmax and Km) of the enzyme was calculated from graphical representations according to the Michaelis and Lineweaver-Burk methods (Hans Lineweaver, 1934).

#### 2.7.12. Data analysis

Data analyses were performed using SAS software version 9 (Inc. Cary NC USA). The experiments were carried out in triplicate. Analysis of variance (ANOVA) and means comparisons were done by Duncan’s multiple range tests.

## 3. Results

### 3.1. Cultural profile used for enzyme production

Figure 1 shows the cultivation profile of aspartic protease produced by *Aspergillus oryzae* DRDFS13 under SSF for 6 days using WB media. The fungus was able to grow in stationary conditions and secrete the enzyme to the culture media starting from the 4^th^ fermentation day with the highest milk-clotting activity (175 U/mL) and protease activity (231 U/mL) recorded on the 5^th^ and 6^th^ days of fermentation time, respectively. It can also be seen that after the 5^th^ fermentation day, the milk-clotting activity decreased slightly. Since, the highest ratio for MCA/PA was detected on the 5^th^ fermentation day; the crude enzyme extract from the 5^th^ fermentation day was used for enzyme purification.

**Figure 1:**
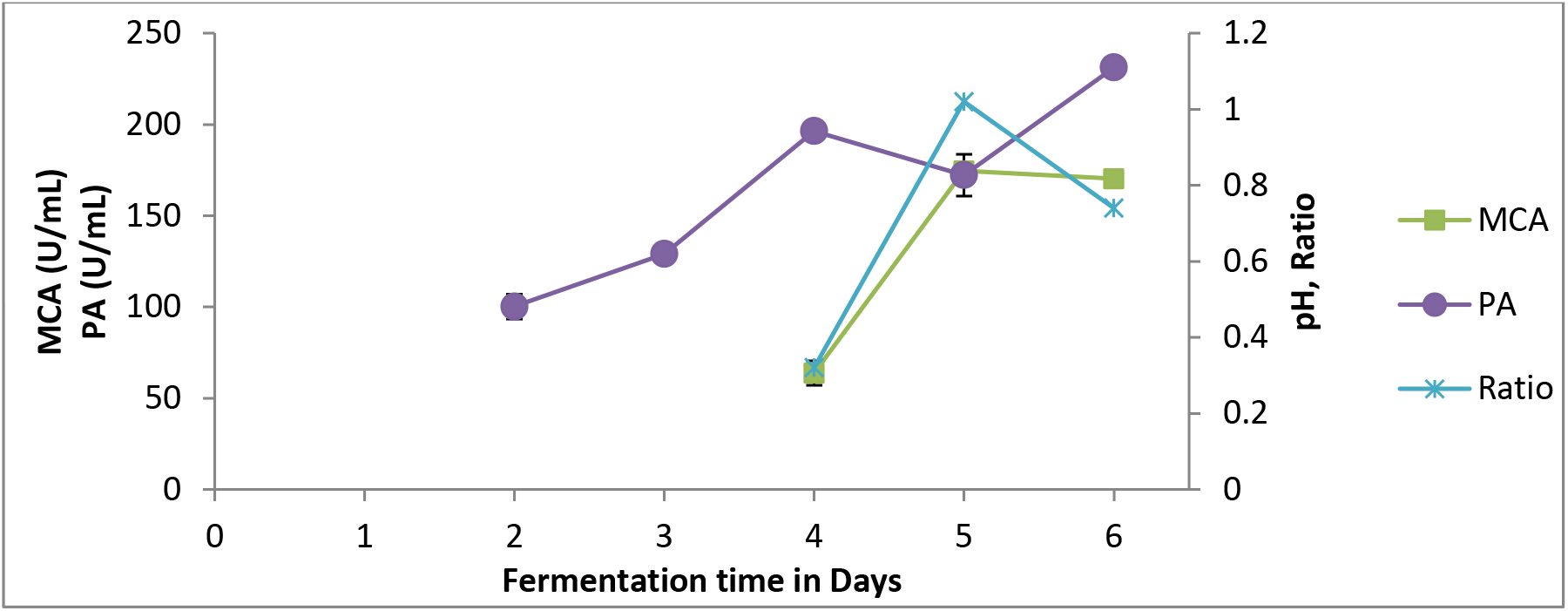
pH, MCA, PA and ratio (MCA/PA) of the crude enzyme extract from *A. oryzae* DRDFS13

### 3.2. Recovery and purification of aspartic protease

The crude enzyme extract from *A. oryzae* DRDFS13 exhibiting a milk-clotting activity (174.61 U/mL) was separated by chromatography using DEAE-sepharose column (IEC) and sepharose 12 pre-packed column (SEC) and eluted with stepwise NaCl gradient. The chromatographic profile of IEC is shown in fig. 2. There are several major protein peaks; however, the prominent peak observed at about 155 min and 310 retention volumes were shown significant milk-clotting activity. Further fractionation of the IEC were shown fractions A_7–10_ contained the target enzyme activity (Fig 3.) whereas the highest MCA (477.11 U/mL) was obtained from the active fraction A_8_. After IEC, active fraction A_8_ was purified 6.2-fold from a crude enzyme, with a final yield of 9.2% and an observed specific activity of 183.5 U/mg. The MCA/PA of active fraction A_8_ was also increased by 3.3 fold. However, significant MCA were also detected from SEC (186.86 U/mL), IEC A_7_ (283.01 U/mL), A_9_, (73.89 U/mL) and A_10_ (9.38 U/mL) (Fig. 3 and Table 4).

**Figure 2:**
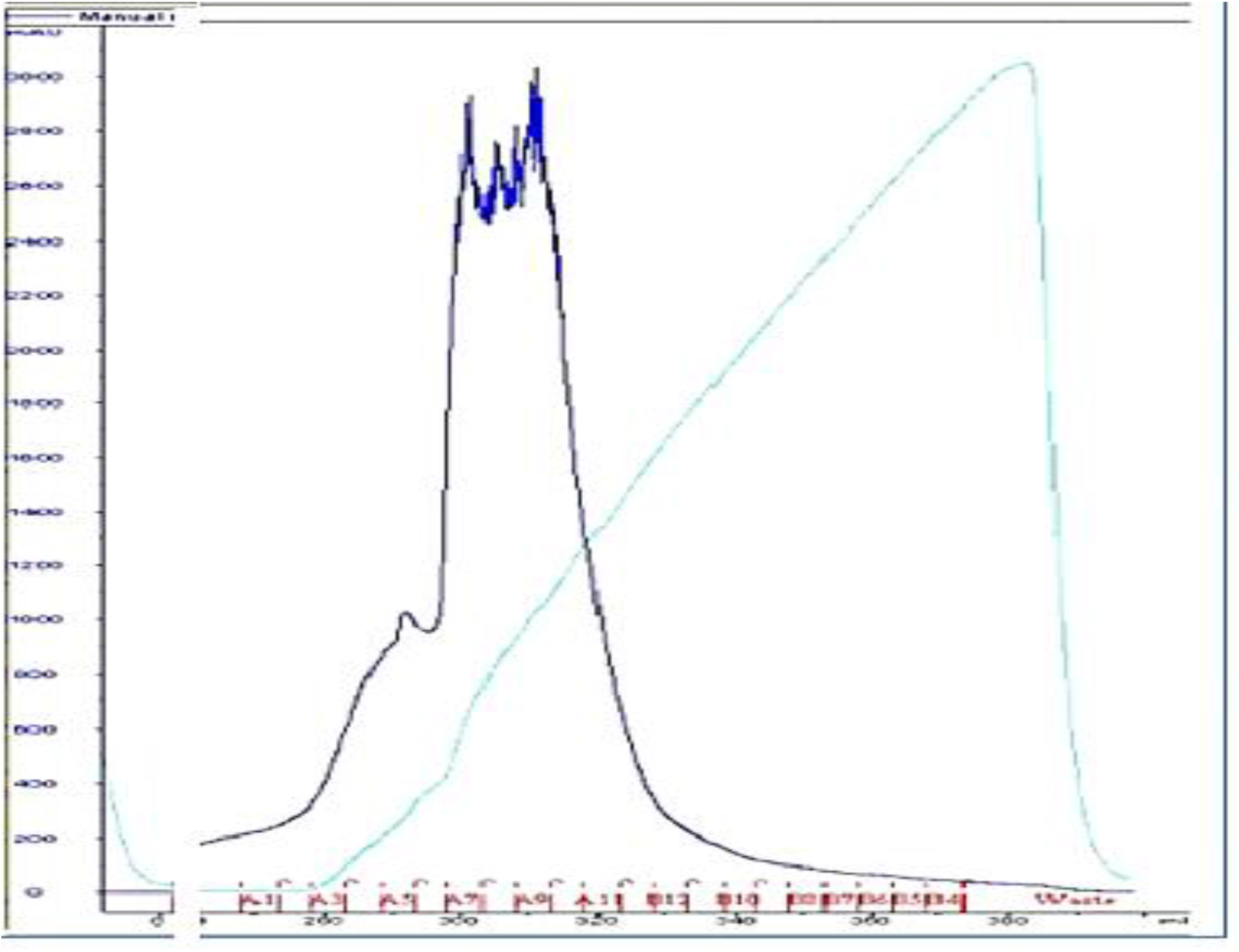
Peaks of ion exchange chromatography obtained after three hours. The x-axis represents retention volume in mL; the Y-axis represents the mAU

**Figure 3:**
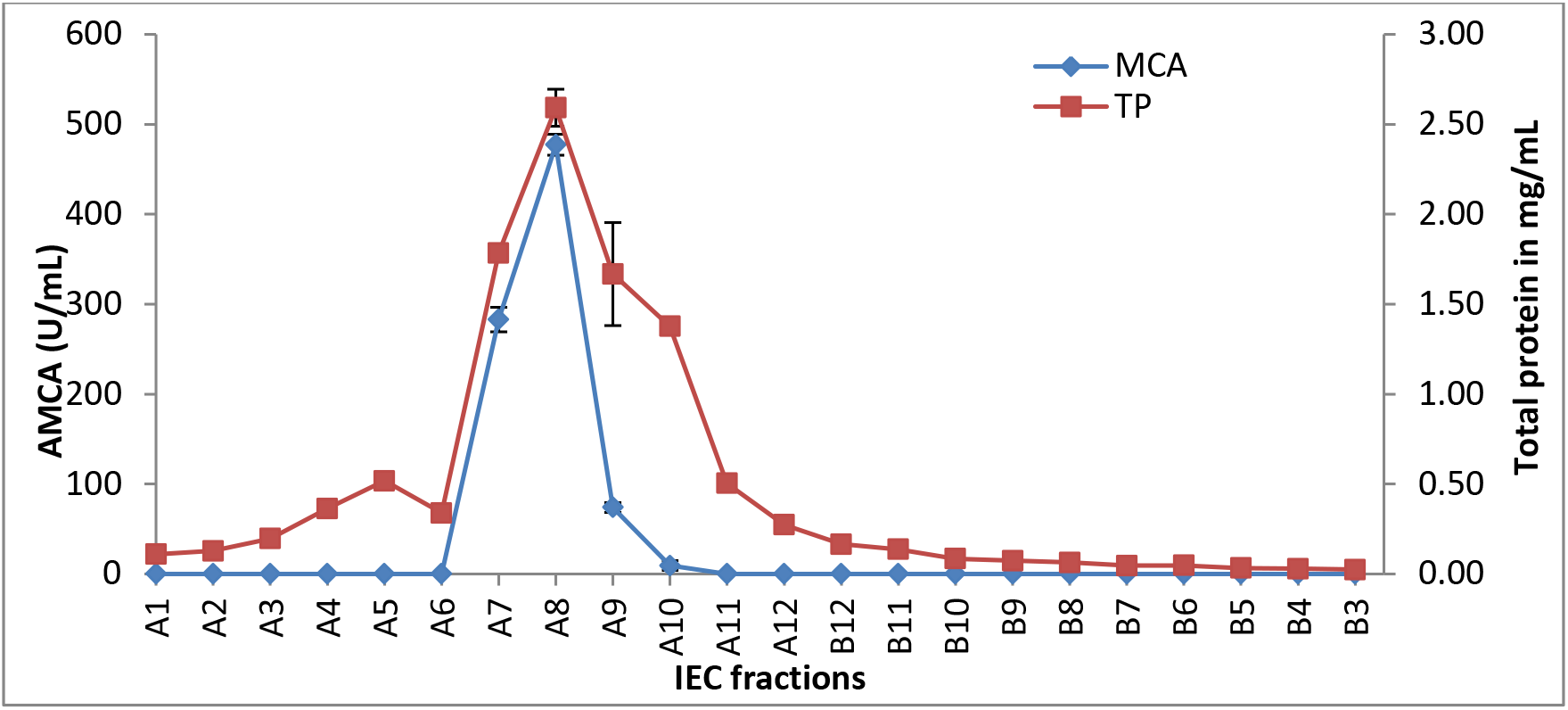
Line graph for MCA and TP of IEC Fractionate from *A. oryzae* DRDFS13 (A_1_-B_3_ is ion-exchange chromatography fractions)

**Table 4:**
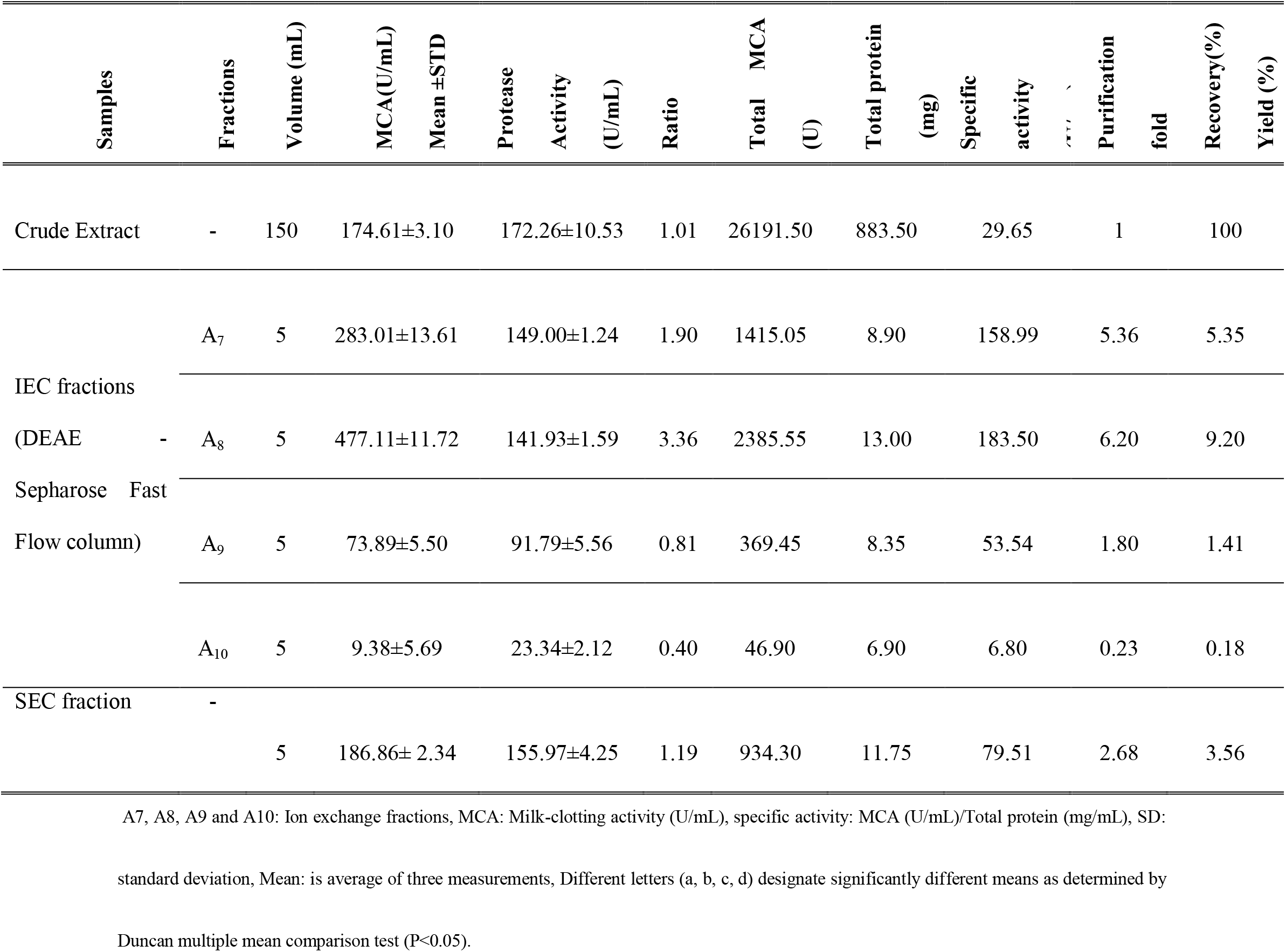
MCA, specific activity and purification fold of aspartic protease from *A. oryzae* DRDFS13 purified by an-ion exchange, size exclusion chromatography

### 3.3. Molecular characterization of purified aspartic protease

Enzyme purity was confirmed by SDS-PAGE and the active IEC fraction A_8_ migrated as a single band with an apparent molecular weight of 40 KDa (Fig 4B, Fig 5A and Table 3). Whereas three bands (one clear band at about 32 kDa mark and 2 small bands at around 11KDa) were obtained from SEC (Fig 4C). Similarly, three bands (one at about 32 kDa marks, the second band at slightly above the 56 kDa marks and the third (two small bands) around the 11 kDa marks) were visualized from crude enzyme on SDS-PAGE gel (Fig 4A).

**Figure 4:**
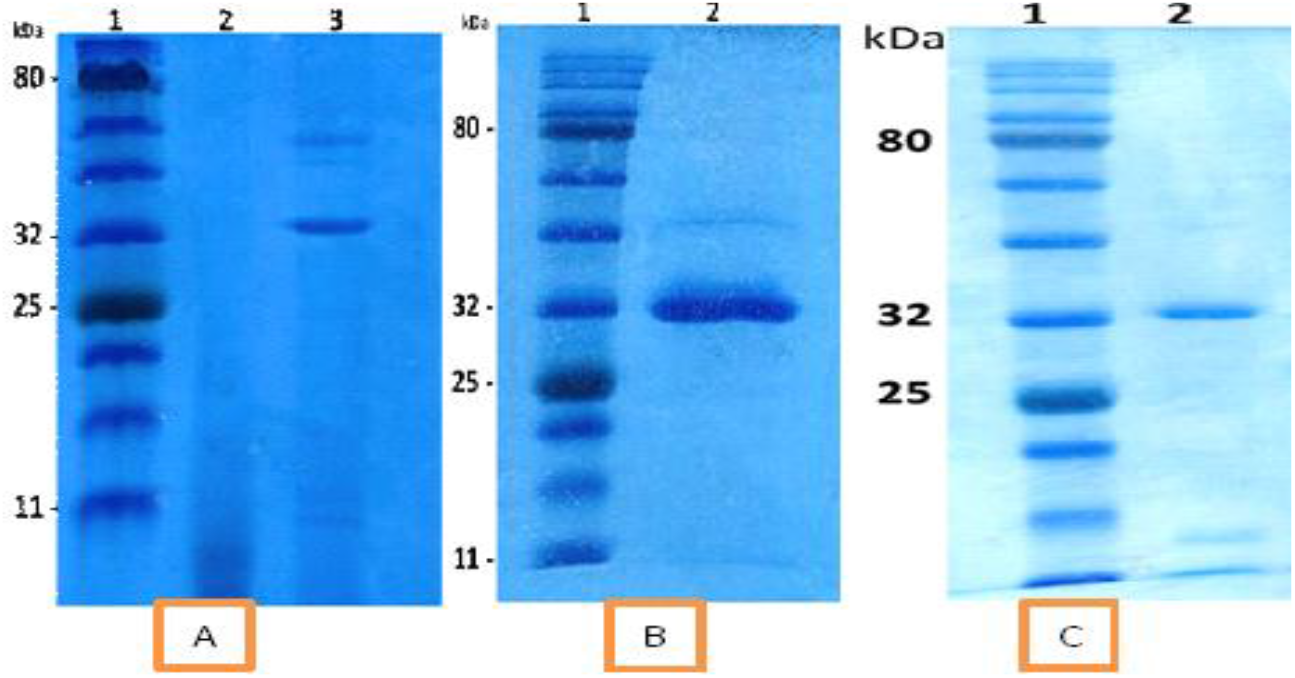
(A) depicts the SDS-PAGE for the crude enzyme extract, well-1: protein marker with pre-stained color plus, well-2: negative control, well-3: crude enzyme extract. (B) depicts the SDS-PAGE for IEC fraction A_8_: well-1: protein marker, well-2: IEC fraction A_8_ (C) depicts the SDS-PAGE for pure enzyme after size exclusion chromatography: well-1: protein marker, well-2: SEC fraction.

**Figure 5:**
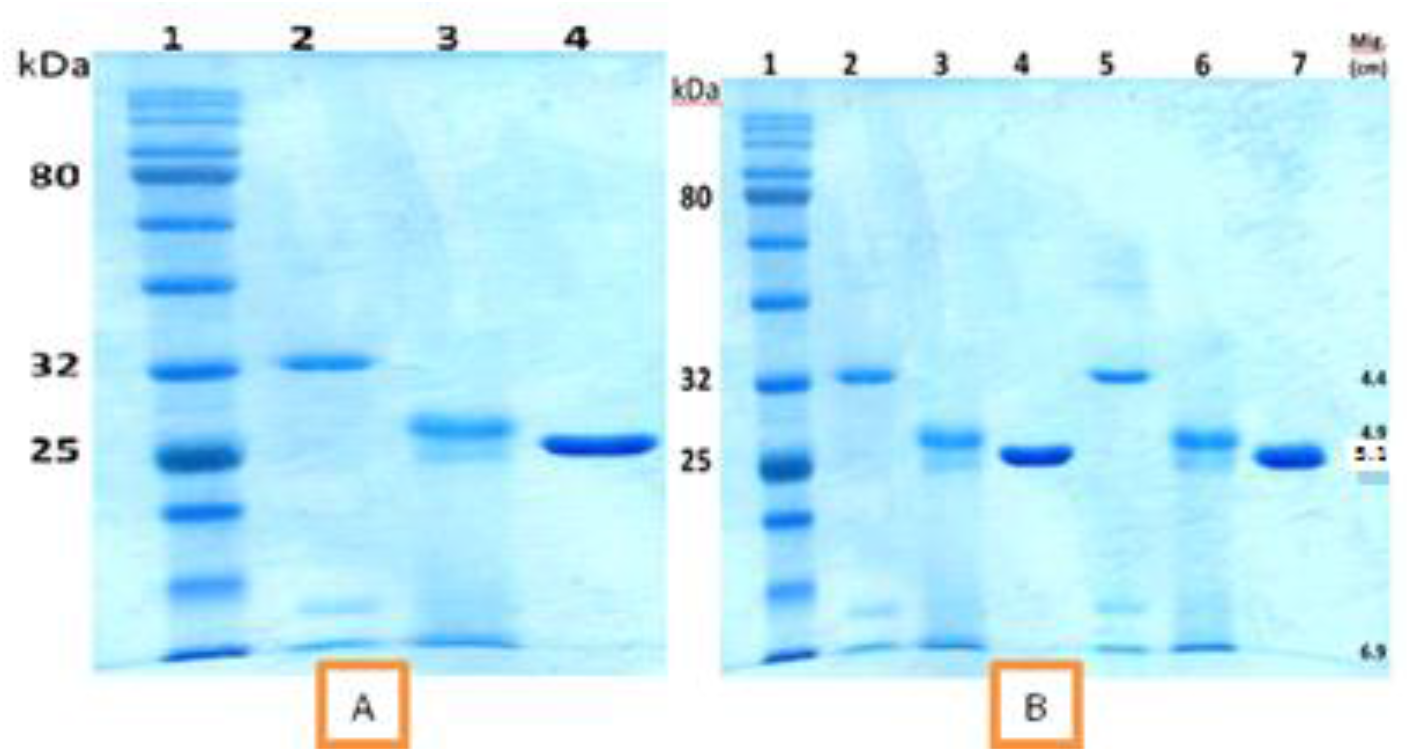
(A) depicts SDS-PAGE for the purified enzyme (A_8_) before and after protein de-glycosylation and Endo-H. Well 1 contains the protein marker, 2 contain the pure enzyme (non-deglycosylated), and 3 contain the protein after de-glycosylation and well 4 contain the pure Endo H (B): depicts the migration distances of the bands in the SDS-PAGE gel Non-de-glycosylated enzyme, glycosylated enzyme, and Eno-H. Well, 1 contains the Protein marker, Wells 2 and 5 contain the pure enzyme, and well 3 and 6 contain the protein after the deglycosylation assay. Lastly, wells 4 and 7 show the pure Endo H enzyme used for the assay.

In this study, de-glycosylation of the aspartic proteinase (IEC A_8_) from *A. oryzae* DRDFS13 was performed by using Endo H. The evidence from SDS-PAGE showed the presence of N-linked protein glycosylation, since a change in molecular weight was observed after treatment with Endo-H indicating the removal of glycan from enzymes (Fig 5A, B). It can be seen that the glycosylated, deglycosylated and Endo-H have a MW of 40, 30 and 27 KDa, respectively (Table 5).

**Figure 6:**
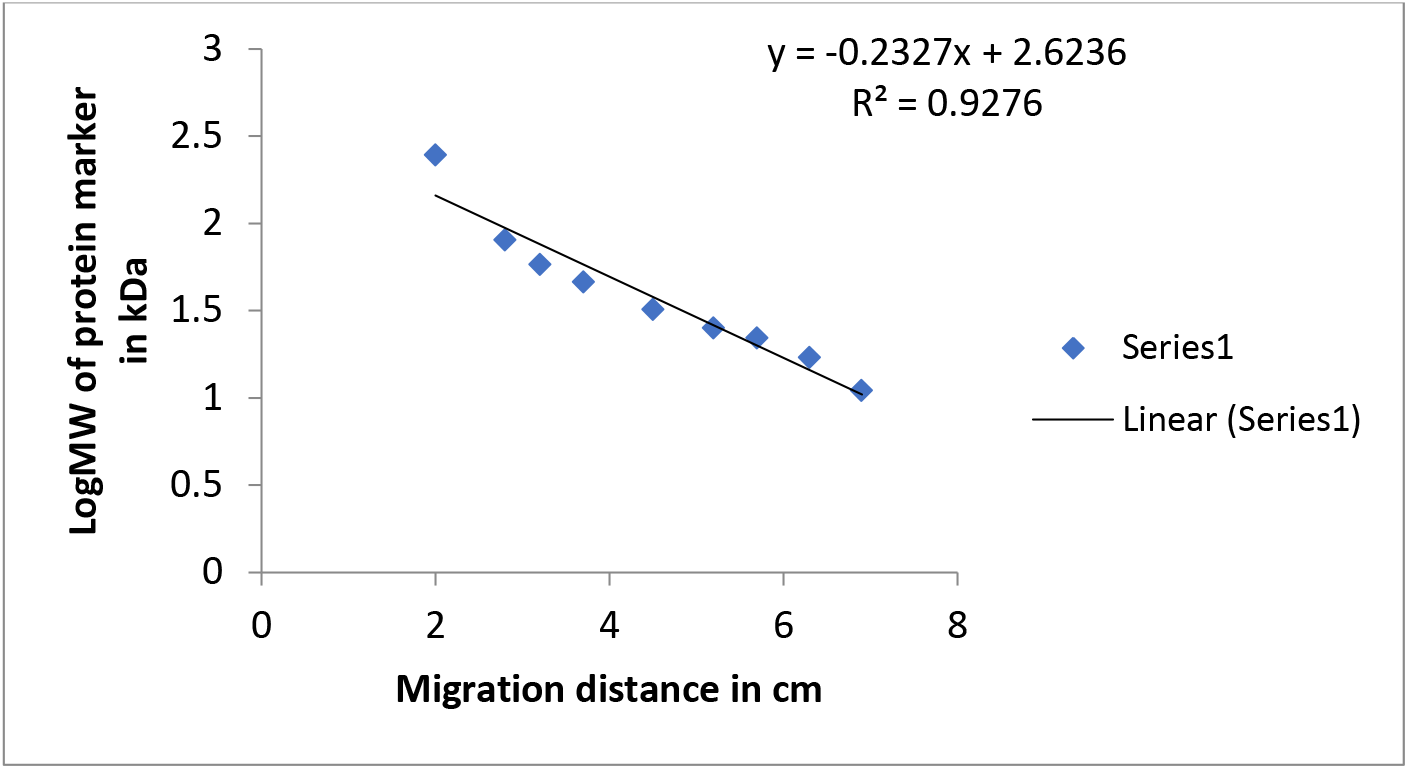
depicts the log MW (y-axis) versus the migration distance in cm (x-axis) of the bands on the marker protein.

**Table 5:** The particular molecular weight assigned for the bands on the marker protein with their respective R_f_ values

| Sample | MW( kDa) | Rf value |
| --- | --- | --- |
| Non-deglycosylated enzyme (2 and 5) | 40 | 0.63 |
| The deglycosylated enzyme (3 and 6) | 30 | 0.71 |
| Endo-H (4 and 7) | 27 | 0.74 |

MW: molecular weight, Rf: relative mobility

The purified proteinase from *A. oryzae* (IEC A_8_) was subjected to inhibition studies with several class-specific agents (Table 6). While iodoacetamide, PMSF and EDTA showed slight effect on milk-clotting activity at concentrations up to 10 mM, pepstatin A caused a substantial inhibition (94 %) on milk-clotting activity of the enzyme.

**Table 6:** Inhibition study of IEC fraction A_8_ from *A. oryzae* DRDFS13

| Inhibitors | Conc | MCA (U/mL) | Residual MCA |
| --- | --- | --- | --- |
| Pepistatin A | 0.02 mM | 116.24±1.41<sup>f</sup> | 24.34 |
|  | 0.04 mM | 55.17±0.13<sup>g</sup> | 11.55 |
|  | 0.06 mM | 46.88±0.09<sup>g</sup> | 9.81 |
|  | 0.08 mM | 33.30±2.05<sup>g</sup> | 6.97 |
|  | 0.1 mM | 28.69±1.57<sup>g</sup> | 6.01 |
| Iodoacetamide | 1 mM | 417.68±10.90<sup>b</sup> | 87.45 |
|  | 10 mM | 396.94±9.84<sup>bc</sup> | 83.11 |
| EDTA | 5mM | 384.22±9.22<sup>cd</sup> | 80.44 |
|  | 10mM | 376.47±23.53<sup>cd</sup> | 78.82 |
| PMSF | 1mM | 360.22±26.88<sup>d</sup> | 75.42 |
|  | 10mM | 318.01±6.32<sup>e</sup> | 66.58 |
| Control (IEC fraction A<sub>8</sub>) | – | 477.62±2.38<sup>a</sup> | 100.00 |

MCA: Milk-clotting activity (U/mL), IEC: Ion-exchange chromatography; pep A: Pepstatin A; EDTA: Ethylene diamine tetraacetic acid; PMSF:
Phenylmethyl fluoride sulphonyl, ND = Not determined, SD: standard deviation, Mean: is average of three measurements, Different letters (a, b,
c, d, e, f, g) designate significantly different means as determined by Duncan multiple mean comparison test (P<0.05).

Due to the shortage of purified enzyme (IEC A_8_), the dialyzed protease was used for further biochemical enzyme characterization such as determination of optimum temperature and temperature stability, determination of optimum pH and pH stability, the effect of cations, effect substrate concentration, and Km and Vmax. Dialysis of the crude enzyme using membrane tube was purified the enzyme 2.2 fold, with a final yield of 5.63% and a detected specific activity of 65.3 U/mg (Table 7).

**Table 7:**
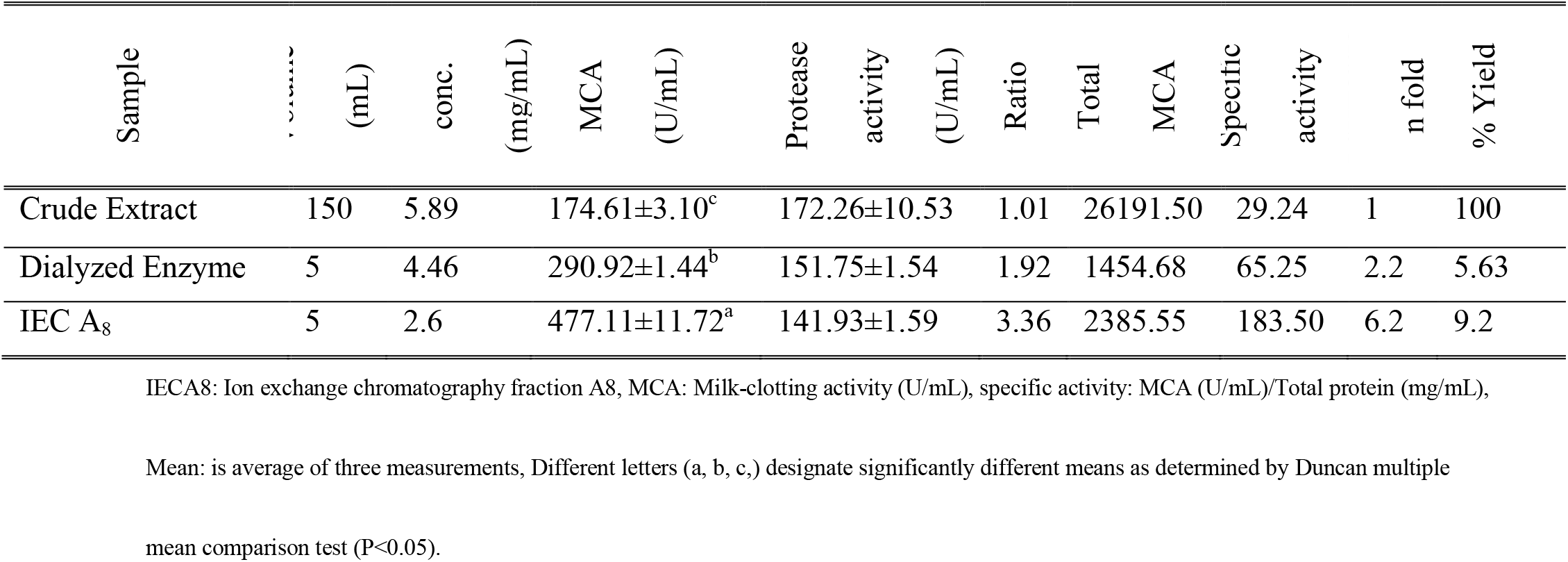
MCA, Specific activity, purification fold, and % yield of aspartic protease enzyme dialyzed and purified by an-Ion exchange chromatography

The catalytic activity of the dialyzed enzyme from *A. oryzae* DRDFS13 was determined at 35 °C and pH 6.5 at varying casein concentrations (0-20 mg/mL). The Km and Vmax of the enzyme was 17.50 mM and 1369 U (Fig 7). The enzyme also showed a good correlation fit of R^2^=0.9862.

**Figure 7:**
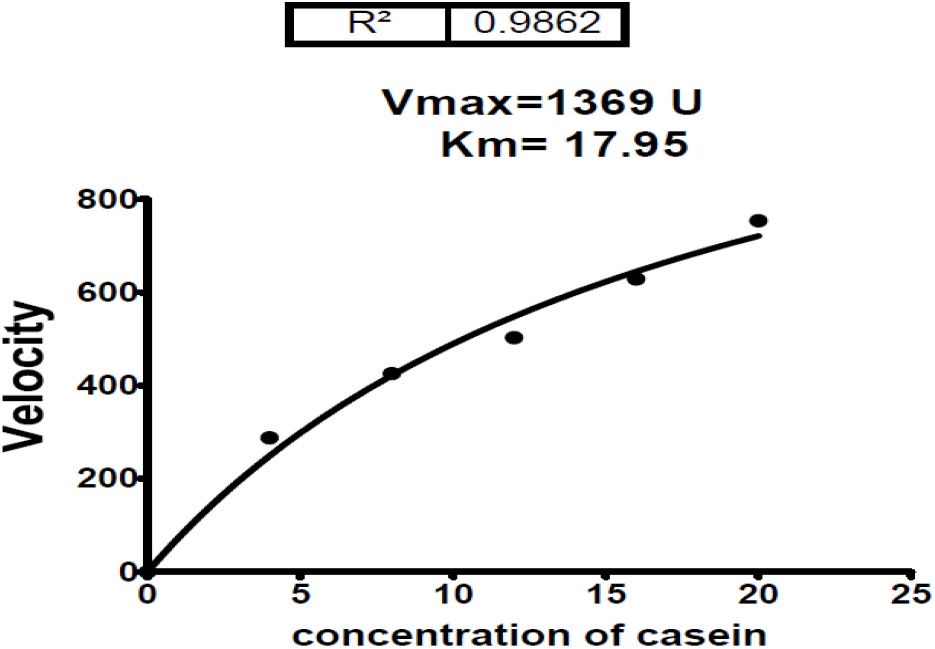
Km and Vmax for the dialyzed enzyme from *A. oryzae* DRDFS13

The relative milk-clotting activity (%) was studied as a function of the pH of the skim-milk substrate at 35 °C (Fig. 8). The dialyzed enzyme was most active between pH values 4.5-6.5 with maximum activity at pH 5.0. Its activity decreased by more than 50% at a pH 7.0 and was insignificant at pH 8.0.

**Figure 8:**
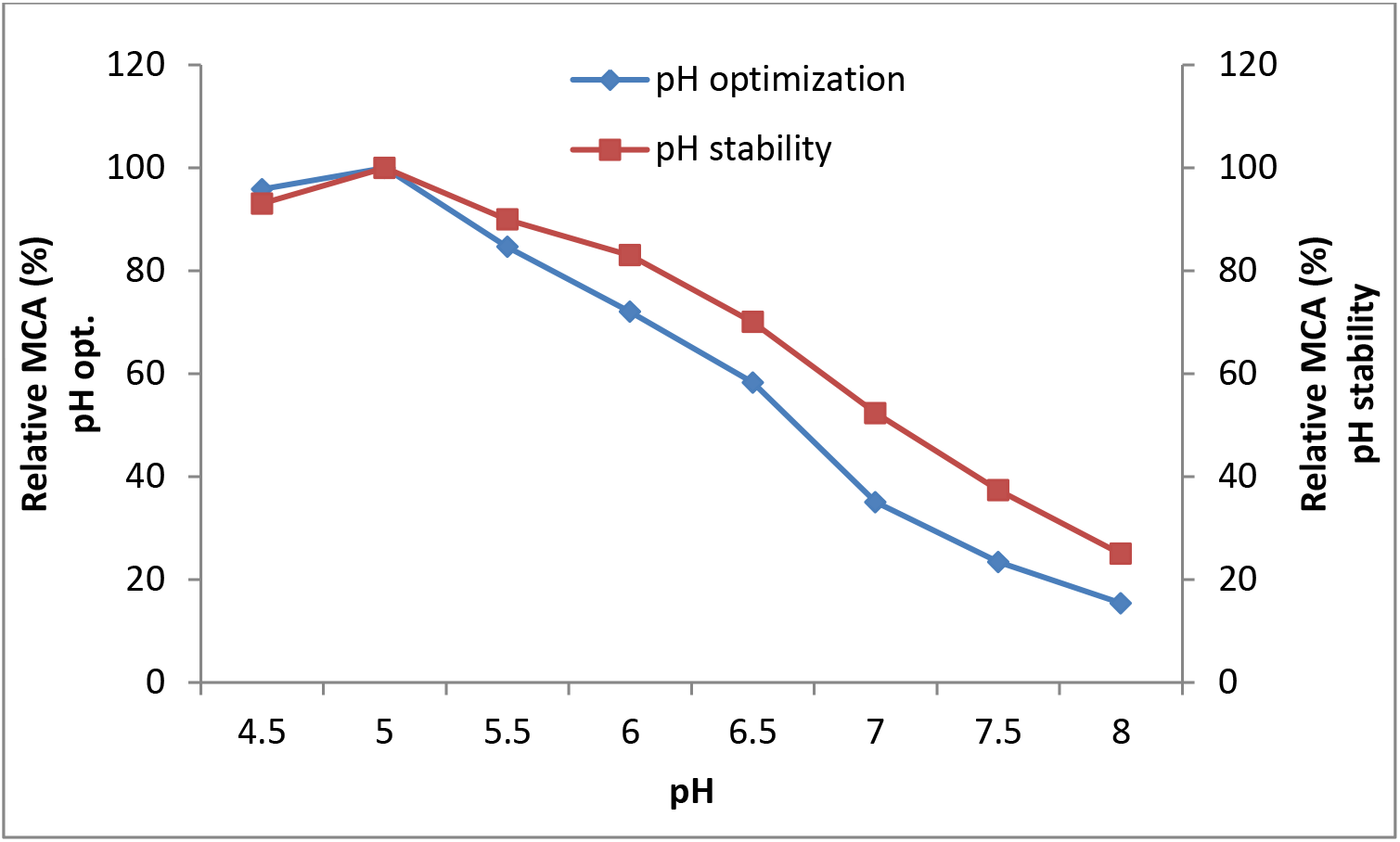
pH optimization and stability test for the dialyzed enzyme from *A. oryzae* DRDFS13

As shown in Fig. 9, the milk-clotting activity of the dialyzed protease from *Aspergillus oryzae* DRDFS13 increased with increasing temperature and reaching a maximum at 60 °C. Beyond 60 °C, the milk-clotting activity of the protease was started to decline sharply.The enzyme retained more than 85% of its MCA at a temperature between 35 to 45 °C (Fig. 10).

**Figure 9:**
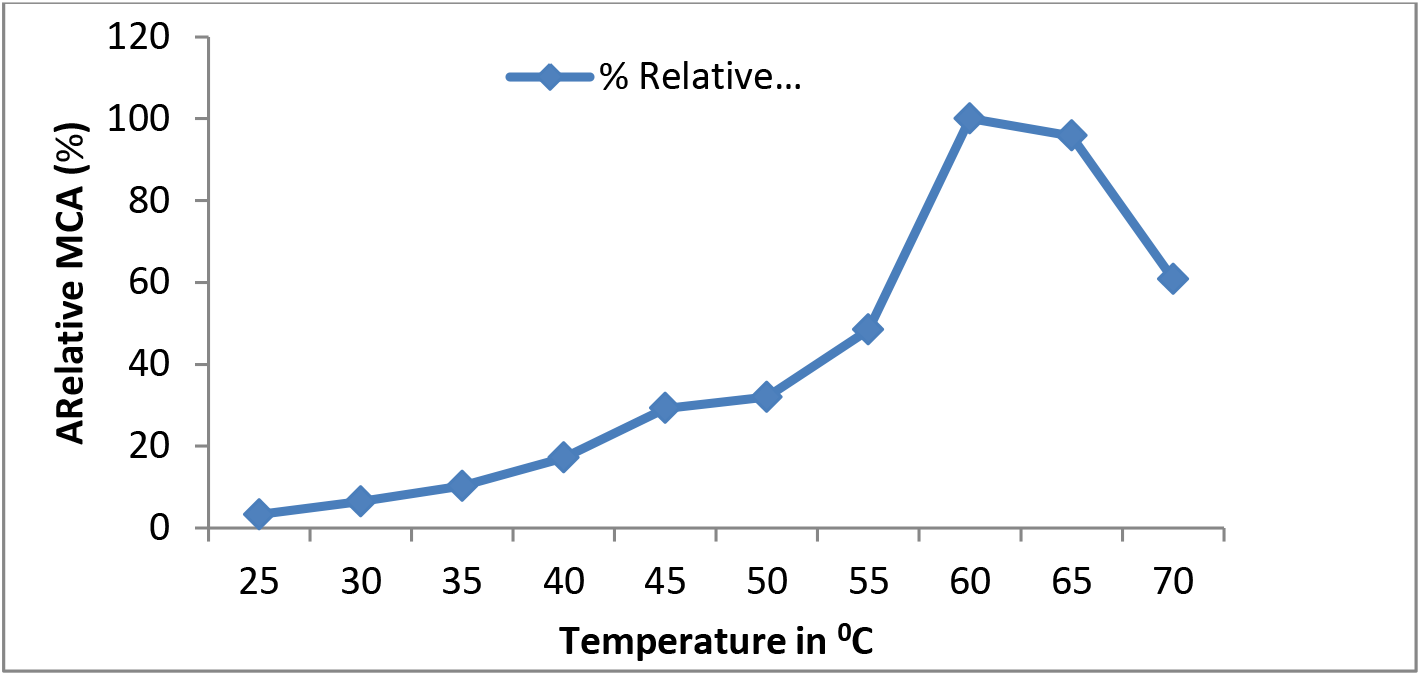
Temperature optimization test for a dialyzed extract from *A. oryzae* DRDFS13

**Figure 10:**
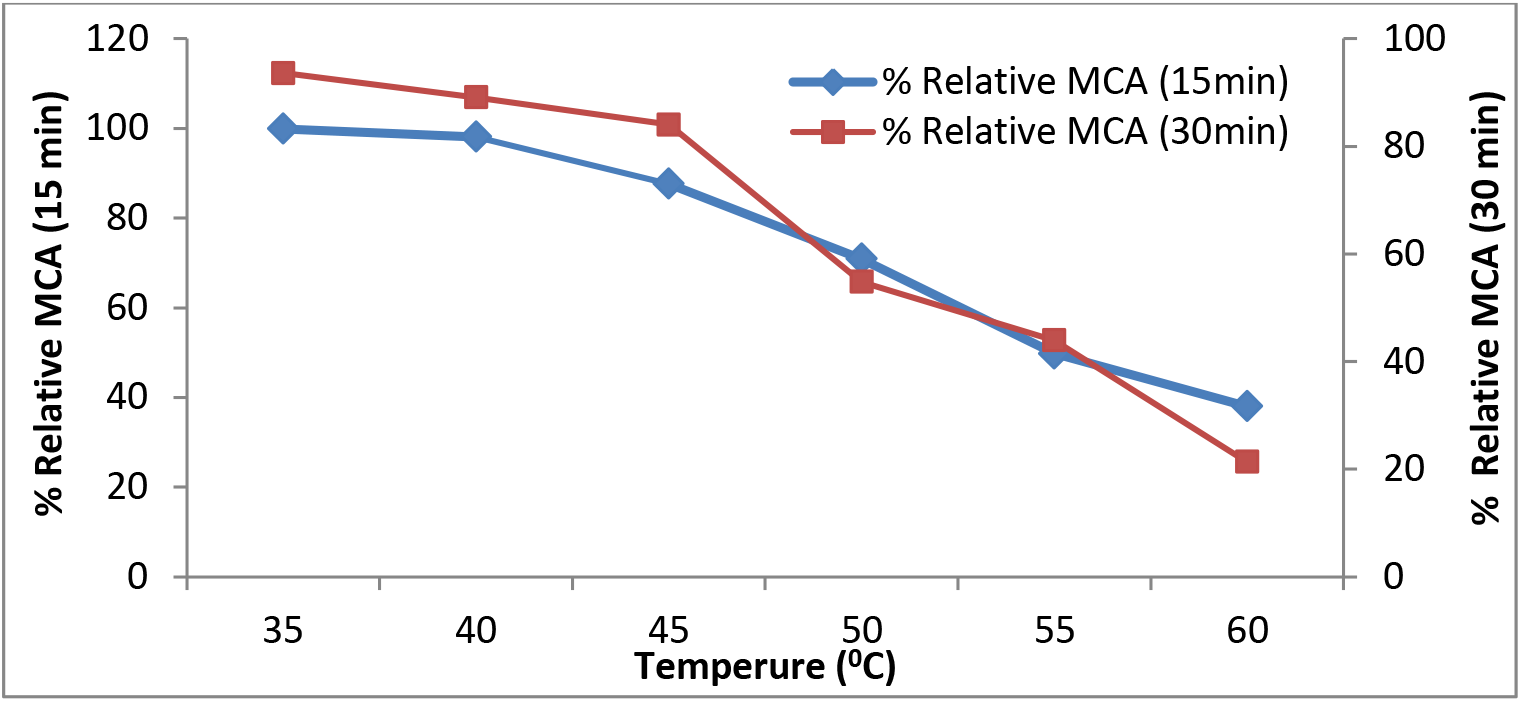
Temperature stability test for a dialyzed extract from *A. oryzae* DRDFS13

The effect of cations (10 mM) on milk-clotting activity was examined at 35 °C using skim-milk as a substrate (Table 8). Accordingly, most cations gave a significant effect on the enzyme activity except NiCl_2_ and ZnSO_4_ that inhibited the MCA as compared to the control. However, the highest MCA (649 U/mL) was obtained from MnSO_4_. Further optimization of MnSO_4_ concentration, showed the highest MCA (1550.00±50.00 U/mL) at 50 mM (Fig.11).

**Table 8:**
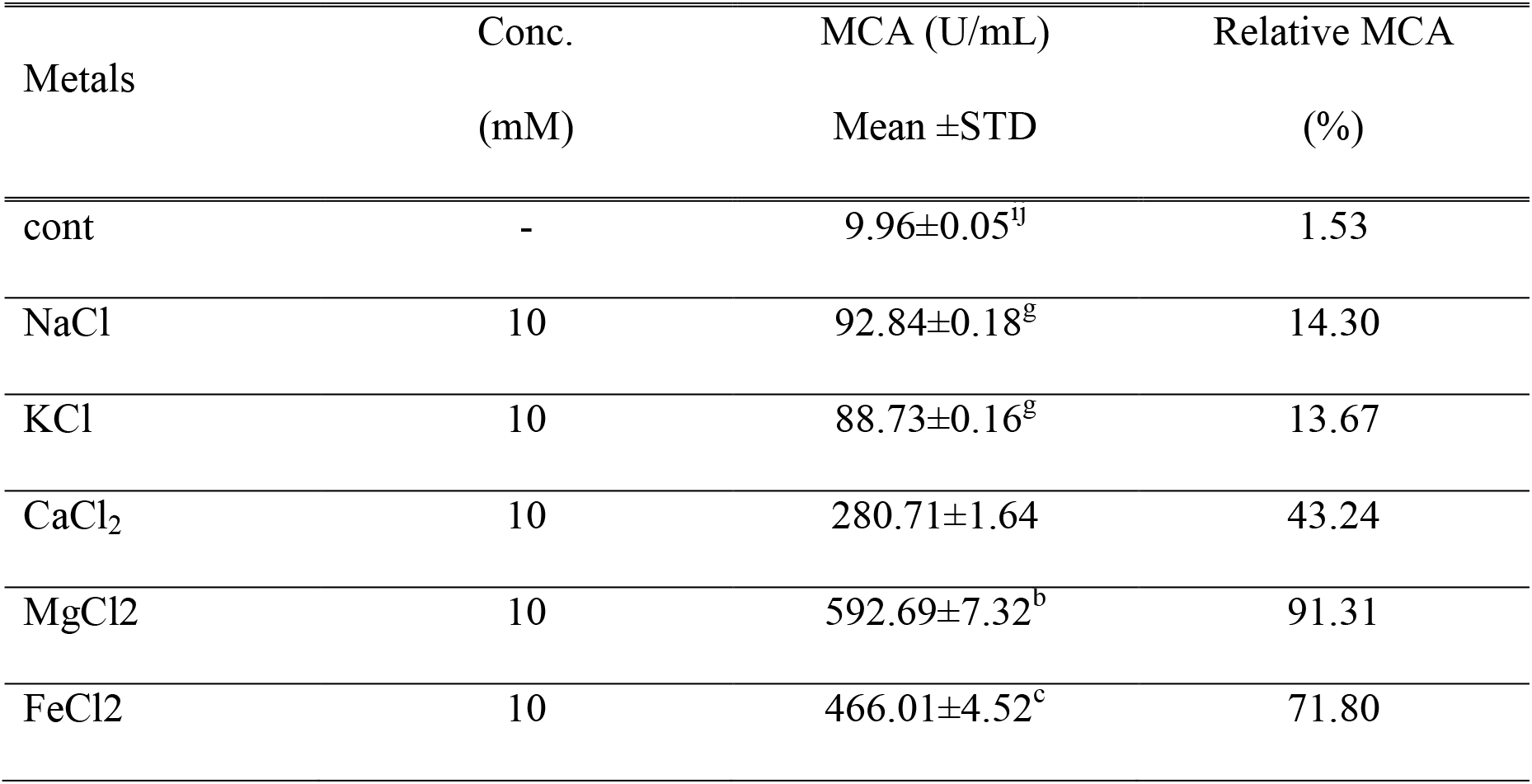
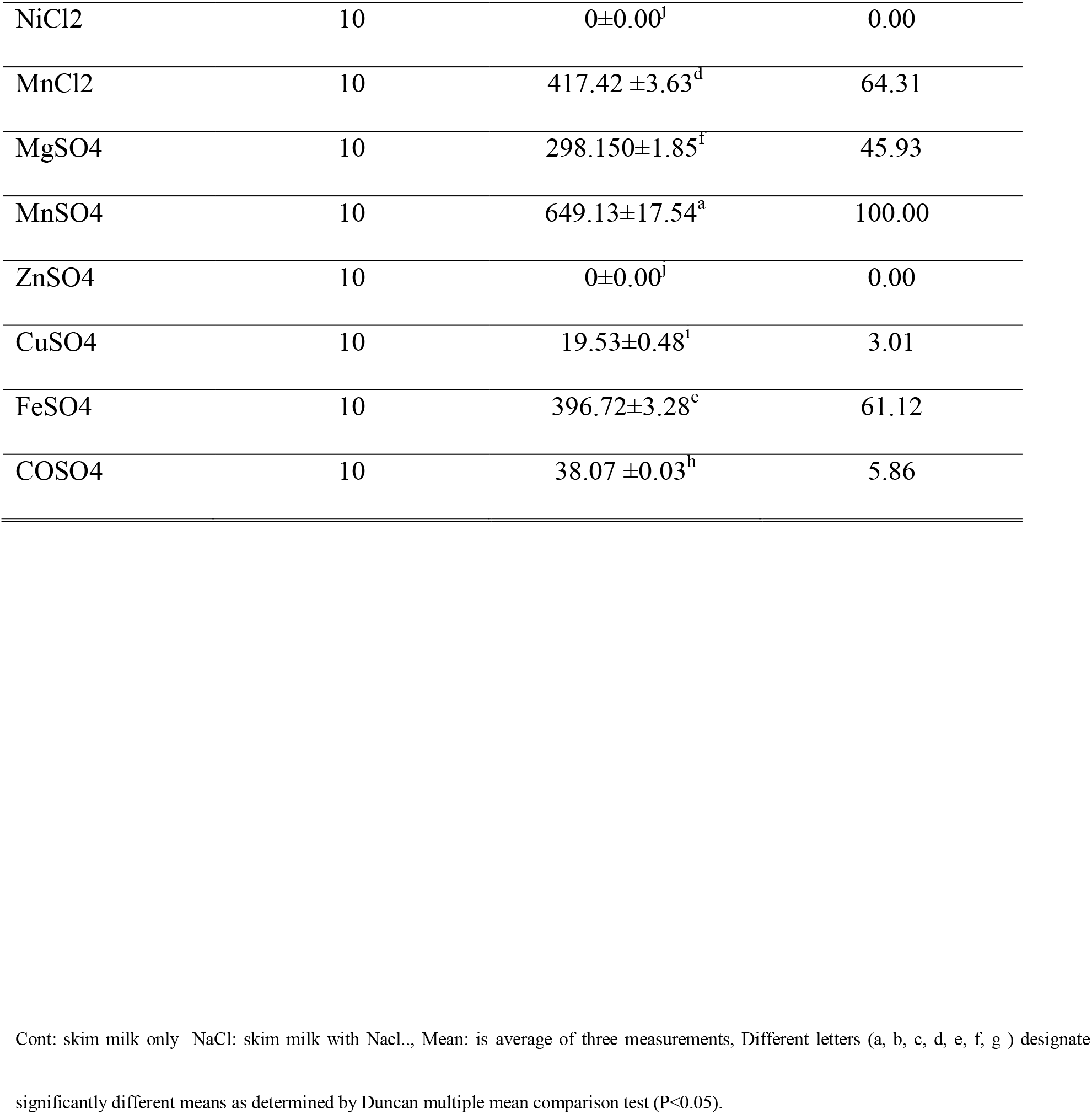
The effect metal ions on MCA of the dialyzed enzyme from *A. oryzae* DRDFS13

| Metals | Conc. | MCA (U/mL) | Relative MCA |
| --- | --- | --- | --- |
| | (mM) | Mean ±STD | (%) |
| cont | - | $9.96 \pm 0.05^{\text{ij}}$ | 1.53 |
| NaCl | 10 | $92.84 \pm 0.18^{\text{g}}$ | 14.30 |
| KCl | 10 | $88.73 \pm 0.16^{\text{g}}$ | 13.67 |
| $\text{CaCl}_2$ | 10 | $280.71 \pm 1.64$ | 43.24 |
| $\text{MgCl}_2$ | 10 | $592.69 \pm 7.32^{\text{b}}$ | 91.31 |
| $\text{FeCl}_2$ | 10 | $466.01 \pm 4.52^{\text{c}}$ | 71.80 |
| NiCl<sub>2</sub> | 10 | 0±0.00<sup>j</sup> | 0.00 |
| MnCl<sub>2</sub> | 10 | 417.42 ±3.63<sup>d</sup> | 64.31 |
| MgSO<sub>4</sub> | 10 | 298.150±1.85<sup>f</sup> | 45.93 |
| MnSO<sub>4</sub> | 10 | 649.13±17.54<sup>a</sup> | 100.00 |
| ZnSO<sub>4</sub> | 10 | 0±0.00<sup>j</sup> | 0.00 |
| CuSO<sub>4</sub> | 10 | 19.53±0.48<sup>i</sup> | 3.01 |
| FeSO<sub>4</sub> | 10 | 396.72±3.28<sup>e</sup> | 61.12 |
| CoSO<sub>4</sub> | 10 | 38.07 ±0.03<sup>h</sup> | 5.86 |

Cont: skim milk only NaCl: skim milk with NaCl., Mean: is average of three measurements, Different letters (a, b, c, d, e, f, g ) designate
significantly different means as determined by Duncan multiple mean comparison test (P<0.05).

**Figure 11:**
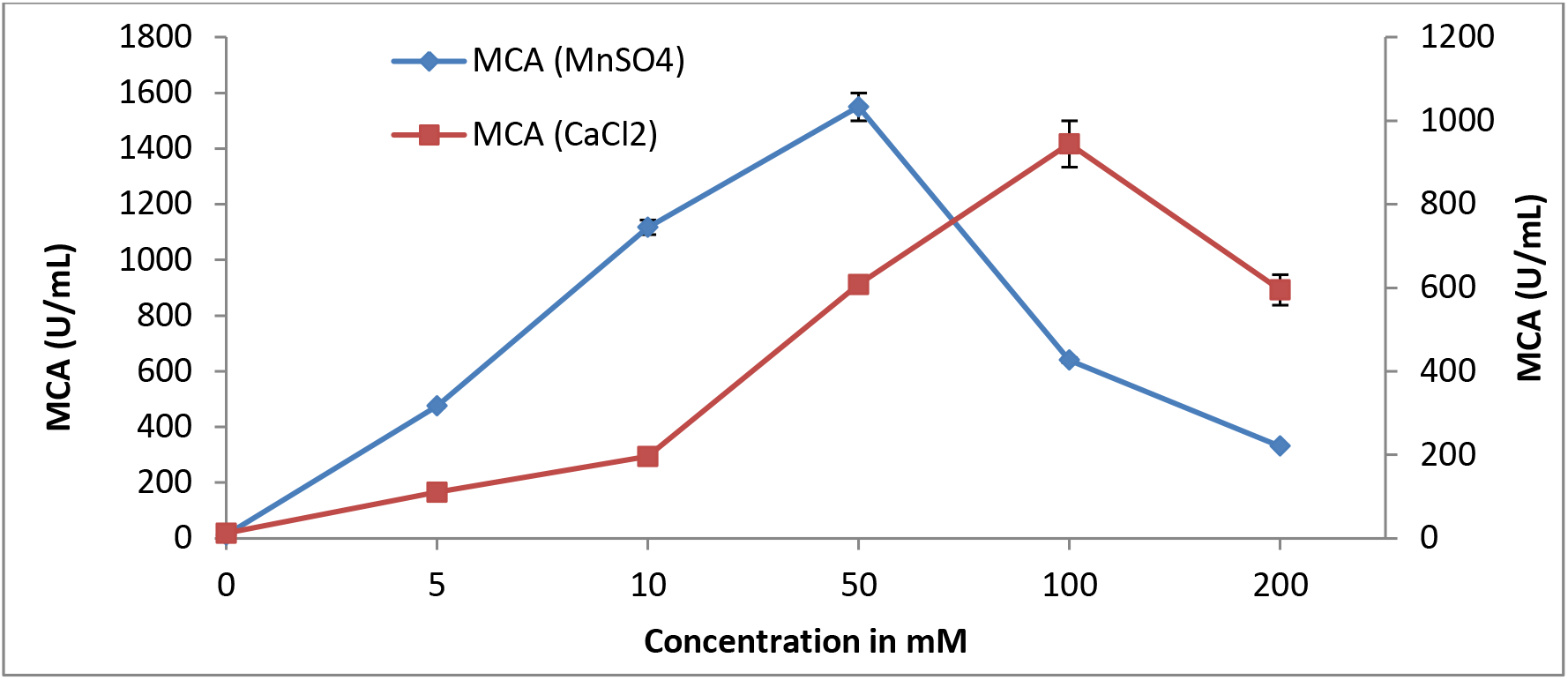
The effect of MnSO_4_ and Cacl_2_ concentration on MCA of the enzyme from *A. oryzae* DRDFS13

The present data also showed substrate concentration had an effect on the milk-clotting activity of the dialyzed enzyme in that the highest MCA (905.98±17.09 U/mL) was obtained at 200 g/L skim-milk concentration (Table 9).

**Table 9:** The effect of skim milk conc. on MCA of the enzyme from *A. oryzae* DRDFS13

| Conc. of Skim milk | MCA (U/mL) | Relative MCA (%) |
| --- | --- | --- |
| 25 | $107.41 \pm 1.68^e$ | 11.86 |
| 50 | $161.63 \pm 1.63^d$ | 17.84 |
| 100 | $246.32 \pm 6.32^c$ | 27.19 |
| 150 | $539.39 \pm 6.06^b$ | 59.54 |
| 200 | $905.98 \pm 17.09^a$ | 100.00 |

MCA: Milk-clotting activity (U/mL), SD: standard deviation, Mean: is an average of three measurements, Different letters (a, b, c, d) designate significantly different means as determined by Duncan multiple mean comparison test ( $P < 0.05$ ).

## 4. Discussion

In this study, we purified an extracellular aspartic protease from *Aspergillus oryzae* DRDFS13 by means ion exchange and size-exclusion chromatography and investigate its biochemical characteristics using 10% solution of skim-milk.

The highest milk-clotting activity (175 U/mL) and MCA/PA ratio (1.02) of the crude enzyme from *A. oryzae* DRDFS13 were observed at 120 h of incubation time at 30 °C under SSF. However, the biomass levels reached maximum (19.77±8.20 mg/gdfs) at 48 h and decreased afterwards. This implies that the fungus secrete the enzyme after its maximum growth is attained by utilizing various carbon and nitrogen sources in solid-substrate. Similar to the present study, a milk-clotting activity between 46.4 to 160.3 U/mL (Merheb-Dini et al. 2010), 60.5 CU/mL (Silva *et al*., 2014) and 165 U/mL (Mukhtar 2015) were obtained from the crude enzyme extracted from *Thermomucor indicae-seudaticae* N31, *Thermomucor indicae-seudaticae* N3 and *M. pusillus* IHS6, respectively.

Purification of the enzyme by dialysis, SEC and IEC revealed that, the active fractions A_8_ obtained after IEC was purified 6.2-fold from a crude enzyme, with a yield of 9.2%, milk-clotting activity of 477 U/mL and an observed specific activity of 183.5 U/mg. The MCA/PA of the same IEC fraction was also increased by 3.3 fold. The present finding suggests that ion-exchange chromatography is an appropriate purification step in improving the activity of fungal aspartic protease as compared to simple dialysis and size-exclusion chromatography.

Similar to the present study, aspartic proteases purified from *A. oryzae* BCRC 30118 (Yin, Chou, and Shann-Tzong Jiang 2013) and *Phanerochaete chrysosporium* (Rodrigues et al. 2017) showed high enzyme activity with a specific activity of about 117.62 U/mg and 224.7 U/mg, respectively. The purification fold of 10.0 (Ao *et al*., 2018) and 9.0 (Fazouane-Naimi et al. 2010) gained for the neutral protease from *A. oryzae* Y1 after DEAE-Sepharose Fast Flow chromatography and for aspartic protease enzyme from *A. niger* FFB1after gel filtration was comparable with the present study. However, the specific activity of 1.2 U/mg (Nouani *et al*., 2011) and (248.1 U/g) (Souza et al. 2017) obtained for an aspartic protease from *M. pusillus* and *A. foetidus*, respectively after gel filtration were smaller than this study. The differences recorded in the specific activity and purification fold from this study and others could be due to variation in microbes and/or enzyme purification methods used.

Enzyme purity was also confirmed by SDS-PAGE and the active IEC fraction A_8_ migrated as a single band with a molecular weight of 40 KDa. This value was in the range of molecular weights normally determined for aspartyl proteinases (32-45 kDa) (Hsiao *et al*., 2014). The molecular weight of the purified enzyme was similar with MW reported for aspartic protease from *Rhizopus oryzae* (≈39 KDa) (Hsiao *et al*., 2014), *Phanerochaete chrysosporium* (38 kDa) (Rodrigues et al. 2017), *A. oryzae* BCRC 30118 (41 KDa) (Yin et al. 2013). However, the molecular weight of the purified enzyme was higher than reported for aspartic protease from *M. mucedo* DSM 809 (33 kDa) (Yegin *et al*., 2012) and *Mucor species* (33.5 KDa) (Fernandez-Lahore et al. 1999), *A. oryzae* (32 kDa)(Dhurway *et al*., 2012). The difference in MW could be due to varies type of aspartic protease can be produced from different microorganisms.

The data from de-glycosylation assay exhibited the reduction of molecular weight of the purified protein from 40 KDa to 30 KDa. This indicates the presence of N-linked protein glycosylation as the change in the molecular mass was observed after treatment with Endo-H. This post-translational modification of the aspartic protease enzyme increased its thermal stability and resistant to proteolysis (Goettig 2016; Wang et al. 2008). Similar to this study, two potential N-glycosylation sites were identified for the amino acid sequence of purified recombinant aspartic protease from *Pichia pastori*s (Sun et al. 2018). The purified enzyme produced in this study was better in thermal stability and resistance to proteolysis than the aspartyl proteinase from *M. mucedo* that lacks of N-linked protein glycosylation (Yegin *et al*., 2012).

The purified aspartic protease (IEC A_8_) from *A. oryzae* DRDFS 13 was strongly inhibited by pepstatin A, with nearly complete inactivation happening at an inhibitor concentration of 0.1 mM. Similar results were reported by Hsiao *et al*., (2014); Merheb-Dini *et al*., (2010); Rodrigues *et al*., (2017); Souza *et al*., (2017); Yin *et al*., (2013). In contrast, other inhibitors (Iodoacetamide, PMSF, and EDTA), did not show significant inhibition even at higher concentration of 10 mM. The nearly complete inhibition could be due to specific and irreversible binding of pepstatin A within the active site of aspartate and hence it protects the enzyme-substrate (skim-milk) binding. Strong inhibition by pepstatin A (aspartic protease inhibitor), suggests that the purified protein (IEC A_8_) is classified as an aspartic protease (Hsiao *et al*., 2014).

The results of the dialyzed enzyme from *Aspergillus oryzae* DRDFS13 showed an apparent Km and Vmax value of 17.95 mg/mL (1.80%) and 1369 (U/mL) μmol/mL/min (22.82*10^-3^mmol/mL/sec), respectively. The catalytic result from the present study suggested a wide specificity of the enzyme towards casein substrates and requires further determination of the Km and Vmax using purified enzyme to increase the enzyme affinity towards substrate and increase its reaction velocity. The Km and Vmax value estimated for protease from *A. oryzae* (4.9 mg/mL and 5446.3 U/g) (Janser *et al*., 2014) and acid protease from *Onopordum acanthium* (12.25 mM and 1329.6 U/mL) (Benkahoul *et al*., 2016) using casein substrate was similar to the present study. The Km (19.5 mg/mL) value recorded for the purified MCE from *M. pusillus* QM 436 using casein as a substrate was also comparable with this study (El-Tanboly *et al*., 2013).

On the other hand, the Km and Vmax value estimated for purified milk-clotting enzyme from *A. candidus* (0.059 mg/mL and 8.59*10^-3^ mmol/ml/sec) (Baskar *et al*., 2014), for acidic protease from *A. oryzae* using hemoglobin as a substrate (0.12 mM, 14.29 μmol/min) (Yin et al. 2013) and for acid protease from *A. foetidus* for azocasein substrate (1.92 mg/mL; 0.8 mM, 357.14 U/mL) (Souza *et al*., 2015) were different from the present study. The observed variation in Km and Vmax value between the present and previous study could be due to a dialyzed enzyme (not a purified enzyme) and/or different substrates used for the determination of catalytic activity.

The dialyzed enzyme from *A. oryzae* DRDFS13 was shown best milk-clotting activity between pH values 4.5-6.5 with highest activity at pH 5.0. This implies that the enzyme is active at acidic pH and appropriate for cheese production (Kumari et al. 2016). Similarly, the optimal milk-clotting activity for aspartyl proteinase from *Mucor mucedo* DSM 809 (Yegin *et al*., 2012), milk-clotting enzyme from *Rhizomucor miehei* (Moghaddaml *et al*., 2008), acid protease from *A. niger* FFB1 (Fazouane-Naimi *et al*., 2010), milk-clotting protease from *Thermomucor indicae-seudaticae* N31 (Silva *et al*., 2014) and milk-clotting enzyme from *Thermomucor indicae-seudaticae* N31 (Merheb-Dini *et al*., 2010) were observed at pH 5.0–5.6, 5.3, 5.5, 5.6 and 5.7, respectively. On the other hand the highest milk-clotting activity for milk-clotting enzyme from *A. oryzae* MTCC 5341(Vishwanatha *et al*., 2010) and milk-clotting enzymes from *A. flavo furcatis* (Alecrim *et al*., 2015) were detected at pH 6-6.3 and 7, respectively.

Regarding pH stability, the dialyzed enzyme retained (83-100%) of milk-clotting activity during 1h incubation at pH range from 4.5-6.0. However, its activity decreased by more than 50% at pH 7.0 and was insignificant at pH 8.0. Therefore, the stability of the enzyme in MCA in the present study corresponds with a milk pH (5.0 to 6.5) at the time of enzyme addition that is appropriate for cheese production (Kumari et al. 2016). The instability of the enzyme at alkaline pH’s suggested an irreversible inactivation of enzyme (Kumari et al. 2016) or alkaline pH’s may cause a protein denaturation in the dialyzed enzyme (Fazouane-Naimi et al. 2010). Similar stability values were also found for proteinase from *M. mucedo* (pH 5.0–5.5) (Yegin *et al*., (2012), milk-clotting protease from *Thermomucor indicae-seudaticae* N31(pH 3.5-6.0) (Merheb-Dini *et al*., 2010), rennet like enzyme from *A. niger* (pH 3.0-5.5) (Fazouane-Naimi *et al*., 2010) and milk-clotting enzyme from *A. flavo furcates* (pH 4.0-6.0) (Alecrim *et al*., 2015). In contrast, the stability values noticed at pH 3.5-4.5 for the milk-clotting protease from *Thermomucor indicae-seudaticae* N31 (Silva *et al*., 2014) and at pH 5.0-8.0 for milk-clotting enzyme from *A. oryzae* MTCC 5341 (Vishwanatha *et al*., 2010) were different from this study.

The maximum milk-clotting activity of the dialyzed enzyme was obtained at 60 °C. The result from this study corresponds with that of Kumari *et al*., (2016) that optimum temperature of majority of MCEs lies in the range of 30-75 °C. Similarly, the optimum temperature observed for the milk-clotting/acid protease activities recorded for milk-clotting protease from *Thermomucor indicae-seudaticae* (65 °C)(Silva *et al*., 2014), aspartic peptidase from *Phanerochaete chrysosporium* (60-65 °C) (Rodrigues et al. 2017), milk-clotting enzyme from *Rhizomucor miehei* (65 °C) (Moghaddaml *et al*., 2008), milk-clotting enzyme from *A. oryzae* MTCC 5341 (55 °C) (Vishwanatha *et al*., 2010) and acid protease enzyme from *A. awmari* (55 °C) (Souza *et al*., 2015) were equivalent to the present study. However, the optimum temperature noticed for milk-clotting/acid protease activities for milk-clotting enzyme from *A.candidus* (35-50 °C) (Baskar *et al*., 2014), milk-clotting protease from *A. niger* FFB1 (45 °C) (Fazouane-Naimi *et al*., 2010), aspartyl proteinase from *M. mucedo* DSM 809 (35-40 °C)(Yegin *et al*., 2012) and milk-clotting enzyme from *A. flavor furcates* (40 °C) (Alecrim *et al*., 2015) were lower than this study.

The thermostability profile of dialyzed enzyme from *A. oryzae* DRDFS13 showed that the enzyme was stable in the temperature range from 35-45 °C by retaining ≥85% of its MCA activity upon 15 min and 30 min incubation. Further incubation of the enzyme at 55 °C for 15 min and 30 min resulted in 50% and 66% loss of its MCA, respectively. Only 21.5% of the MCA of the enzyme was retained after 30 min incubation at 60 °C. This result suggests that aspartic proteases from *A.oryzae* DRDFS13 seem to be stable at temperatures below 45 °C and susceptible to inactivation above this temperature. The nearly complete inactivation of the enzyme activity upon incubation at 55 °C for 30 min may be a technological advantage since no bitterness development can occur during cheese ripening due to the inactivated proteolytic action of the enzyme after the cooking process of the curd (Sousa *et al*., 2001). Similar to this study, the rennet like enzyme from *A. niger* FFB1 (Fazouane-Naimi *et al*., 2010), milk-clotting enzyme from *Aspergillus oryzae* MTCC 5341 (Vishwanatha *et al*., 2010), milk-clotting enzyme from *Thermomucor indicae-seudaticae* N31 (Silva *et al*., 2014) and aspartyl proteinase from *M. mucedo* DSM 809 (Yegin *et al*., 2012) completely lost their milk-clotting activity at 60 °C, 65 °C, 62 °C, 60 °C and 55 °C upon incubation for 30 min, 15 min, 60 min and 10 min, respectively.

The addition of 10 mM of MgCl_2_, FeCl_2_, MnCl_2_, MnSO_4_, FeSO_4_, and CaCl_2_ showed a pronounced effect on the milk-clotting activity of the enzyme reflecting significant effect on second phase of milk clot formation. However, the highest MCA was obtained from MnSO_4_. Monovalent salts such as KCl and NaCl were also shown slight stimulatory effect on milk-clotting activity. On the other hand, NiCl_2_ and ZnSO_4_ showed an inhibitory effect on the milk-clotting activity as compared to the control (Skim-milk only). Similarly, Ca^2+^ (El-Tanboly *et al*., 2013), Na^+^, K^+^, Mn^2+^ and Ca^2+^ (Liu and Huang, 2015) and Ca^2+^, Mg^2+^, Mn^2+^ (Kumari et al. 2016) and Mn, Ca (Ahmed *et al*., 2016) were shown a noticeable effect on enhancing the the activity of milk-clotting enzyme from *M. pusillus* QM 436, acidic protease from *Rhizopus stolonifer*, MCE from *B. subtilis* MTCC 10422 and MCE from *B. stearothermophilus*, respectively. Whereas the addition of Co^2+^, Cd^2+^ and Zn^2+^ were found to have an inhibitory effect on the activity of MCE from *B. subtilis* MTCC 10422 (Kumari et al. 2016). Generally, low pH, higher temperature and increased levels of calcium improved the MCA of the enzyme perhaps due to it promotes the better enzyme-substrate reaction and hence reduce the clotting time (Alecrim *et al*., 2015; El-Tanboly *et al*., 2013).

The highest MCA was observed at 50 mM MnSO_4_ concentrations. In contrast, the optimal MCA for the rennet-like enzyme from *A.niger* FFB1 were obtained at 0.01 M CaCl_2_(Fazouane-Naimi *et al*., 2010). However, acidic protease from *A. oryzae* BCRC 30118 was not affected by any metals at a concentration of 1.0 mM (Yin et al. 2013). The Milk-clotting activity of aspartic protease from *A. oryzae* DRDFS13 was increased uniformly as the concentration of skim-milk increased from 25 to 200 g/L. The highest MCA was achieved at 200 g/L skim-milk concentration.

## Conclusion

The milk-clotting activity, specific activity, purification fold, biochemical characteristics, inhibition study and results from deglycosylation assay obtained from ion-exchange chromatography fraction A_8_ in the present study confirmed that the enzyme is aspartic protease and could be used as a possible candidate to substitute rennet enzyme for cheese production.

## Declaration

### Ethics approval and consent to participate

Non-applicable

### Consent for publication

Non-applicable

### Availability of data and materials

All data generated or analyzed during this study are publicly available and included in this published article.

### Fund

This work was supported by Microbial, Cellular and Molecular Biology, Addis Ababa University.

### Authors’ contributions

JM: Contributes to conception, acquisition and analysis, interpretation of data, and drafted the work. MK: Contributes to conception, acquisition and analysis, interpretation of data, and drafted the work. JFSO: Contributes to conception, acquisition and analysis, interpretation of data, and drafted the work. FA: contribute to conception, acquisition, and interpretation and substantively revised the work. HMFL: contributes to conception, acquisition, interpretation of data, and drafted the work. All authors have approved the submitted version and the modified version that involves the author’s contribution to the study; and have agreed both to be personally accountable for the author’s own contributions and ensure that questions related to the accuracy or integrity of any part of the work, even ones in which the author was not personally involved, are appropriately investigated, resolved and the resolution documented in the literature.

## Acknowledgment

Moreover, we would like to acknowledge the Department of Microbial, Cellular and Molecular Biology of Addis Ababa University and Downstream Processing Laboratory, Department of Life Sciences and Chemistry, Jacobs University, Bremen, Germany for the facilitation of laboratory space and provision of basic facilities.

## References

Ageitos, J. M., J. A. Vallejo, A. B. F. Sestelo, M. Poza, and T. G. Villa. 2007. “Purification and Characterization of a Milk-Clotting Protease from Bacillus Licheniformis Strain USC13.” Journal of Applied Microbiology 103:2205–13.

Ahmed, Samia A., Hala R. Wehaidy, Osama A. Ibrahimb, Salem Abd ElA. Ghani, and El-Hof Mahmoud. 2016. “Novel Milk-Clotting Enzyme from Bacillus Stearothermophilus as a Coagulant in UF-White Soft Cheese.” Biocatalysis and Agricultural Biotechnology 7:241–49.

Alecrim, Mircella M., Rosana A. Palheta, Maria Francisca S. Teixeira, and Ila Maria De A. Oliveira. 2015. “Milk-Clotting Enzymes Produced by Aspergillus Flavo Furcatis Strains on Amazonic Fruit Waste.” International Journal of Food Science and Technology 50, 50:151–57.

Ao, Xiao et al. 2018. “Purification and Characterization of Neutral Protease from Aspergillus Oryzae Y1 Isolated from Naturally Fermented Broad Beans.” AMB Expr 8(96):1–0.

Arima, Kei, Juhyu Yu, and Shintiro Iwasaki. 1970. “Milk-Clotting Enzyme from Mucor Pusillus Var. Lindt.” Methods in Enzymology 19:446–59.

Baskar, G., S. Babitha Merlin, D. V. Sneha, and J. Angeline Vidhula. 2014. “Production and Purification of Fungal Milk Clotting Enzyme from Aspergillus Candidus.” Journal of Chemical Engineering Research Updates 1:29–34.

Benkahoul, Malika, Ines Bellil, Khelifi Douadi, and Mechakra Aicha. 2016. “Physical and Chemical Properties of the Acid Protease from Onopordum Acanthium: Comparison between Electrophoresis and HPLC of Degradation Casein Profiles.” African Journal of Biotechnology 15(9):331–3440.

Dhurway KS, Ramdas Malakar, Tiwari A, Malviya SN and Yadav M. 2012. “Production, Purification and Characterization of Protease from ‘Aspergillus Oryzae.’” Discovery Biotechnology 1(1):33–38.

El-Tanboly, El-Sayed, Mahmoud El-Hofi, Youssef Bahr Youssef, Wahed El-Desoki, and Azza Ismail. 2013. “Utilization of Salt Whey from Egyptian Ras (Cephalotyre) Cheese in Microbial Milk Clottingenzyme Production.” Acta Sci. Pol., Technol. Aliment. 12(1):9–19.

El-Tanboly El-Sayed; El-Hofi Mahmoud; Bahr Youssef Youssef; El-Desoki Wahed;, and Azza Ismail. 2013. “Utilization of Salt Whey from Egyptian Ras (Cephal Otyre) Cheese in Microbial Milk Clotting Enzymes Production.” Acta Sci. Pol., Technol. Aliment. 12(1):9–19.

Fazouane-Naimi, F. et al. 2010. “Characterization and Cheese-Making Properties of Rennet-Like Enzyme Produced by a Local Algerian Isolate of Aspergillus Niger.” Food Biotechnology 24(3):258–69. Retrieved (<Go to ISI>://000281328300004).

Fernandez-Lahore, H. M. et al. 1999. “Purification and Characterization of an Acid Proteinase from Mesophilic Mucor Sp. Solid-State Cultures.” J. Peptide Res. 53:599–605.

Fernandez-Lahore, H. M., ER. Fraile, and O. Cascone. 1998. “Acid Protease Recovery from a Solid-State Fermentation System.” Journal of Biotechnology 62:83–93.

Goettig, Peter. 2016. “Effects of Glycosylation on the Enzymatic Activity and Mechanisms of Proteases.” Int. J. Mol. Sci. 17:1–24.

Hsiao, Nai-wan et al. 2014a. “Electronic Journal of Biotechnology Puri Fi Cation and Characterization of an Aspartic Protease from the Rhizopus Oryzae Protease Extract, Peptidase R.” EJBT 17(2):89–94. Retrieved (http://dx.doi.org/10.1016/j.ejbt.2014.02.002).

Hsiao, Nai-wan et al. 2014b. “Purification and Characterization of an Aspartic Protease from the Rhizopus Oryzae Protease Extract, Peptidase R.” Electronic Journal of Biotechnology 17:89–94. Retrieved (http://dx.doi.org/10.1016/j.ejbt.2014.02.002).

Janser, Ruann, Soares De Castro, and Helia Harumi Sato. 2014. “Protease from Aspergillus Oryzae: Biochemical Characterization and Application as a Potential Biocatalyst for Production of Protein Hydrolysates with Antioxidant Activities.” Journal of Food Processing 2014(http://dx.doi.org/10.1155/2014/372352):1–12.

Kumari, Rajesh, Bharat Bhushan, Ajay Pal, Anil Panwar, and Sarla Malhotra. 2016. “Purification, Physico-Chemico-Kinetic Characterization and Thermal Inactivation Thermodynamics of Milk Clotting Enzyme from Bacillus Subtilis MTCC 10422.” LWT - Food Science and Technology 65:652–60. Retrieved (http://dx.doi.org/10.1016/j.lwt.2015.08.065).

Liu, Na and Huang, Liang. 2015. “Partial Characterization of an Acidic Protease from Rhizopus Stolonifer.” The Open Biotechnology Journal 9:199–203.

Merheb-Dini, Carolina, Eleni Gomes, Mauricío Boscolo, and Roberto da Silva. 2010. “Production and Characterization of a Milk-Clotting Protease in the Crude Enzymatic Extract from the Newly Isolated Thermomucor Indicae-Seudaticae N31 (Milk-Clotting Protease from the Newly Isolated.” Food Chemistry 120(1):87–93. Retrieved (http://dx.doi.org/10.1016/j.foodchem.2009.09.075).

Moghaddaml, Sh Khalil, M. Khaleghian, F. Naderi, and M. Azinand M. Monajjemi. 2008. “Purification and Characterization of Milk Clotting Enzyme Produced by Rhizomucor Rmiehei.” Journal of Physical and Theoretical Chemistry 5(3):149–54.

Mukhtar, Hamid. 2015. “Production of Rennin-like Acid Protease by Mucor Pusillus through Submerged Fermentation.” (June).

Nouani, A., F. Moulti-Mati, S. Belbraouet, and MM. Bellal. 2011. “Purification and Characterization of a Milk-Clotting Protease from Mucor Pusillus: Method Comparison.” African Journal of Biotechnology 10(9):1655–65.

Ramachandran, N. and R. Arutselvi. 2013. “Partial Purification and Characterization of Protease Enzyme from Nomurarea Rileyi.” International Journal of Pharmaceutical Science and Research 4(9):3460–65.

Rao, Mala B., Aparna M. Tanksale, Mohini S. Ghatge, and Vasanti V. Deshpande. 1998. “Molecular and Biotechnological Aspects of Microbial Proteases †.” MIicrobiology and Molecular Biology Reviews 62(3):597–635.

Rodrigues, Ronivaldo et al. 2017. “Biochemical and Milk-Clotting Properties and Mapping of Catalytic Subsites of an Extracellular Aspartic Peptidase from Basidiomycete Fungus Phanerochaete Chrysosporium.” Food Chemistry 225:45–54.

Silva, B. L., F. M. Geraldes, C. S. Murari, E. Gomes, and R. Da-Silva. 2014. “Production and Characterization of a Milk-Clotting Protease Produced in Submerged Fermentation by the Thermophilic Fungus Thermomucor Indicae-Seudaticae N31.” Applied Biochemistry and Biotechnology 172(4):1999–2011.

Sousa M.J.; Ardo Y.; and McSweeney P.L.H. 2001. “Advances in the Study of Proteolysis during Cheese Ripening.” International Dairy Journal 11(327-345):327–45.

Souza, Paula Monteiro et al. 2015. “Macromolecules Kinetic and Thermodynamic Studies of a Novel Acid Protease from Aspergillus Foetidus.” International Journal of Biological Macromolecules 81:17–21. Retrieved (http://dx.doi.org/10.1016/j.ijbiomac.2015.07.043).

Souza, Paula Monteiro et al. 2017. “Production, Purification and Characterization of an Aspartic Protease from Aspergillus Foetidus.” Food and Chemical Toxicology 109:1103–10.

Souza, Paula Monteiro De et al. 2015. “A Biotechnology Perspective of Fungal Proteases.” Brazilian Journal of Microbiology 46(2):337–46.

Sun, Qian et al. 2018. “A Novel Aspartic Protease from Rhizomucor Miehei Expressed in Pichia Pastoris and Its Application on Meat Tenderization and Preparation of Turtle Peptides.” Food Chemistry 245(October 2017):570–77. Retrieved (https://doi.org/10.1016/j.foodchem.2017.10.113).

Vishwanatha, KS., Rao, A. A. G., and Singh, S. A. 2010. “Production and Characterization of a Milk-Clotting Enzyme from Aspergillus Oryzae MTCC 5341.” Appl Microbiol Biotechnol 85:1849–59.

Vishwanatha, K. S., A. G. Appu and Rao, and Singh Sridevi Annapurna. 2009. “Characterisation of Acid Protease Expressed from Aspergillus Oryzae MTCC 5341.” Food Chemistry 114(2):402–7. Retrieved (http://dx.doi.org/10.1016/j.foodchem.2008.09.070).

Walker, John M. 1996. “The Bicinchoninic Acid (BCA) Assay for Protein Quantitation.” Pp. 11–14 in The Protein Protocols Handbook, 2nd Edition, edited by J. M. Walker. ToTowa: Humana Press Inc.

Wang, Wei et al. 2008. “pH Dependent Effect of Glycosylation on Protein Stability.” European Journal of Pharmaceutical Sciences 33:120–27.

Yegin;, Sirma, Yekta Goksungur;, and Marcelo Fernandez-Lahore; 2012. “Purification, Structural Characterization, and Technological Properties of an Aspartyl Proteinase from Submerged Cultures of Mucor Mucedo DSM 809.” Food Chemistry 133(4):1312–19. Retrieved (http://dx.doi.org/10.1016/j.foodchem.2012.01.075).

Yegin, Sirma, Yekta Goksungur, and Marcelo Fernandez-Lahore. 2012. “Purification, Structural Characterization, and Technological Properties of an Aspartyl Proteinase from Submerged Cultures of Mucor Mucedo DSM 809.” Food Chemistry 133(4):1312–19. Retrieved (http://dx.doi.org/10.1016/j.foodchem.2012.01.075).

Yegin S.; Goksungur Y.; and Fernandez-Lahore M. 2012. “Purification, Structural Characterization, and Technological Properties of an Aspartyl Proteinase from Submerged Cultures of Mucor Mucedo DSM 809.” Food Chemistry 133:1312–1319.

Yin, Li-Jung, Ya-Hui Chou, and Shann-Tzong Jiang. 2013. “Purification and Characterization of Acidic Protease from Aspergillus Oryzae BCRC 30118.” Journal of Marine Science and Technology 21(1):105–10.

